# Novel low-avidity glypican-3 specific CARTs resist exhaustion and mediate durable antitumor effects against hepatocellular carcinoma

**DOI:** 10.1101/2021.07.09.451642

**Authors:** Leidy D Caraballo Galva, Xiaotao Jiang, Mohamed S Hussein, Huajun Zhang, Rui Mao, Pierce Brody, Xiangyang Chi, Yibing Peng, Aiwu Ruth He, Mercy Kehinde-Ige, Ramses Sadek, Xiangguo Qiu, Huidong Shi, Yukai He

## Abstract

Chimeric antigen receptor engineered T cells (CARTs) are being developed to treat solid tumors including hepatocellular carcinoma (HCC). However, thus far, CARTs have not been as effective against solid tumors as compared to blood cancers. A main reason is that, once infiltrating into a solid tumor mass, CARTs are surrounded and chronically stimulated by persistent target antigens, which may drive them to exhaustion. We hypothesize that, due to weak engagement, low-avidity CARTs will resist the antigen-driven exhaustion and apoptosis and maintain effector functions in solid tumors, generating durable antitumor effects. To test this idea, we developed a novel human glypican-3 (hGPC3) specific antibody (8F8) that binds an epitope close to that of GC33 (the frequently used high-affinity antibody), but with ~17 folds lower affinity. *In vitro*, the low-avidity 8F8 CART killed tumor cells and produced effector cytokines to the same extent as high-avidity GC33 CART. Remarkably, however, 8F8 CART expanded and persisted to a greater extent than GC33 CART *in vivo*, resulting in durable responses against HCC xenografts. Compared to GC33 CARTs, there were significantly more (5 times) 8F8-BBz CART detected in the tumor mass. Importantly, the tumor infiltrating 8F8 CARTs were less apoptotic and more resistant to exhaustion, revealed by their enhanced and durable effector functions overtime. We predict that this novel low-avidity 8F8-BBz CART has a greater potential than mainstream high-avidity CARTs in effectively treating patients with HCC or other hGPC3+ solid tumors.

## Introduction

Liver cancer, majority being hepatocellular carcinoma (HCC), is the 6^th^ most common cancer and the 3^rd^ leading cause of cancer death in the world (1), underscoring the urgent need to develop novel therapies. Immunotherapy has become a mainstream cancer treatment, which relies on the antigen-specific T cells that do not always exist and thrive in the tumor (2). One way to provide tumor-specific T cells is to redirect patient’s T cells with T cell receptors (TCR) (3, 4) or chimeric antigen receptors (CAR) (5). The success of CAR modified T cells (CARTs) in blood cancers has sparked tremendous effort to develop CARTs for solid tumors, including HCC (6). Different from blood cancer, solid tumor mass creates a physical barrier to T cell tumor infiltration. Furthermore, after T cells manage to infiltrate into tumor lesion, they submerge into a tumor antigen swamp with complexed stroma matrix that imped T cell movement but also constantly stimulate them without break, driving them into exhaustion and even death (7, 8). Approaches have been studied to reduce CART exhaustion, such as modification of the IgG1 Fc spacer (9), utilization of 4-1BB co-stimulatory domain instead of CD28 (10), and combination with checkpoint blockade (11). However, despite intensive effort, CART therapy has not yet proved as effective for solid tumors as it is for blood cancers (12, 13).

The affinity (strength between a single pair of molecules) of antigen-binding domain and its target has a significant impact on CART’s avidity (total strength of multiple pairs of molecules) and the ensuing activation and antitumor effects (6, 14). Conventionally, high-affinity mAbs are preferred because they induce strong activation and can detect low antigens (15–17). For example, the FMC antibody in CD19 CAR is high affinity (K_D_=0.32nM) (18). The CARs (19–21) targeting human glypican-3 (hGPC3), an oncofetal protein re-expressed in HCC and squamous lung cancers (22), are from high affinity mAbs too, i.e., GC33 (EC_50_=0.24nM) (23), YP7 (EC_50_=0.3nM) (24), and 32A9 (EC_50_=1.24nM) (21). The Phase I clinical trial of GC33 CART generated modest antitumor efficacy at best (25). High-avidity CARTs maybe more prone to activation-induced exhaustion and death due to strong engagement with antigen, especially in the presence of persistent antigens in tumor mass. Recently, several studies suggest that low-avidity CARs can be more beneficial. Low-avidity CART can distinguish antigen high tumor from antigen low normal cells to avoid off-tumor toxicity (17, 26, 27). Report showed that mutations in LFA-1, the ligand for ICAM1 receptor, created a low-avidity CAR with reduced toxicity and better effects (28). In addition, recent study found that low-avidity CD19 CART generated enhanced expansion and prolonged persistence in blood cancers (18). However, another study reported that low-avidity engagement did not prevent anti-GD2 CARTs from being depleted after antigen exposure (29). It remains unknown whether low-avidity CARTs can generate stronger antitumor effects against solid tumors.

In this study, we developed novel low-avidity hGPC3 CARTs and studied their antitumor effect against HCC xenografts. We first generated 3 hGPC3-specific mAbs (6G11, 8F8, and 12D7) with distinct epitopes and affinity. 8F8 mAb binds to an epitope close to that bound by GC33, but has ~17 times lower affinity. The 8F8 CART’s avidity for hGPC3 was 4-5 times lower than GC33 CART. However, the low-avidity engagement was sufficient for 8F8 CART to kill both hGPC3^hi^ HepG2 and hGPC^lo^ Huh7 cells and to produce cytokines to the same extent as high-avidity GC33 CART. Importantly, 8F8 CART generated enhanced *in vivo* expansion, persistence, and durable antitumor effects in treating HCC xenografts. Analysis of CARTs in tumor mass showed that the 8F8-BBz CART, had 5 times more infiltration, was less apoptotic and exhausted, and maintained better function for a longer time. *In vitro* co-culture with tumor cells also showed that 8F8 CARTs were less apoptotic and exhausted, and contained higher central memory T cells than GC33 CARTs. In conclusion, the novel low-avidity hGPC3-specific 8F8 CARTs resist exhaustion and maintain better function in tumor lesions, and thus will likely generate potent therapeutic effects in treating HCC and other GPC3+ solid tumors.

## Materials and Methods (Details in Supplementary Materials and Methods)

### Cells

Details of human T cells isolation and cell cultures were in Supplementary Materials and Methods.

### Mice and tumor models

The use of mice and tumor models were approved by IACUC of Augusta University. Details were in Supplementary Materials and Methods

### hGPC3 specific mAbs

The novel mAbs were generated as previously descried (30) and their cDNA sequences were identified as previously described (3, 31). Recombinant mAbs were produced. Details were in Supplementary Materials and Methods.

### Epitope and affinity of mAbs

The mAb’s epitopes were mapped. The affinity of mAbs to hGPC3 was determined by Biolayer interferometry. Details were in Supplementary Materials and Methods.

### Generation of CARTs

The second generation CARs were created. Human primary T cells were transduced by lentiviral vectors as described (3, 32, 33) to create CARTs. Details were in Supplementary Materials and Methods.

### Adoptive cell transfer

CARTs (10-12 days) of the indicated numbers were injected into tumor-bearing mice via tail vein.

### Immunological analysis

Immune staining, immunological assay, and LDH assay were detailed in Supplementary Materials and Methods

### Tumor-Infiltrating Lymphocytes (TILs)

CARTs in the tumor lesions and their exhaustion and apoptosis status and function were studied.

Details were in Supplementary Materials and Methods.

### Immunohistochemical (IHC) staining

Details of IHC and the scoring were in Supplementary Materials and Methods.

### Bulk RNA Sequencing

CARTs were sorted from tumor lesions and bulk RNAseq was done. Details were in Supplementary Materials and Methods.

### Statistical analysis

Statistical analyses were described on the figures.

## Results

### Novel hGPC3-specific mAbs are developed and characterized

First, we developed hGPC3 mAbs that specifically stained HCC cells and tissues. Hybridomas generated using the splenocytes of hGPC3-immunized mice were screened by ELISA. 22 mAbs were found to bind hGPC3 (Fig.1A). Among them, 14 mAbs stained HepG2 cells (Fig.1B and fig.S1) and 7 of the 14 also stained hGPC3^lo^ Huh7 cells albeit with lower intensity (fig.S2). Since hGPC3 might be naturally cleaved into hGPC3-N (AA_1-358_) and hGPC3-C (AA_359-580_) fragments, we examined the new mAbs bound to which fragment. To this end, the 293T cells transfected with hGPC3-N or hGPC3-C (fig.S3A) were intracellularly stained with the mAbs. The data show that, out of the 14 mAbs that stained HepG2 cells, 11 bound to the hGPC3-N and 3 bound to the hGPC3-C (fig.S3B).

**Fig. 1.**
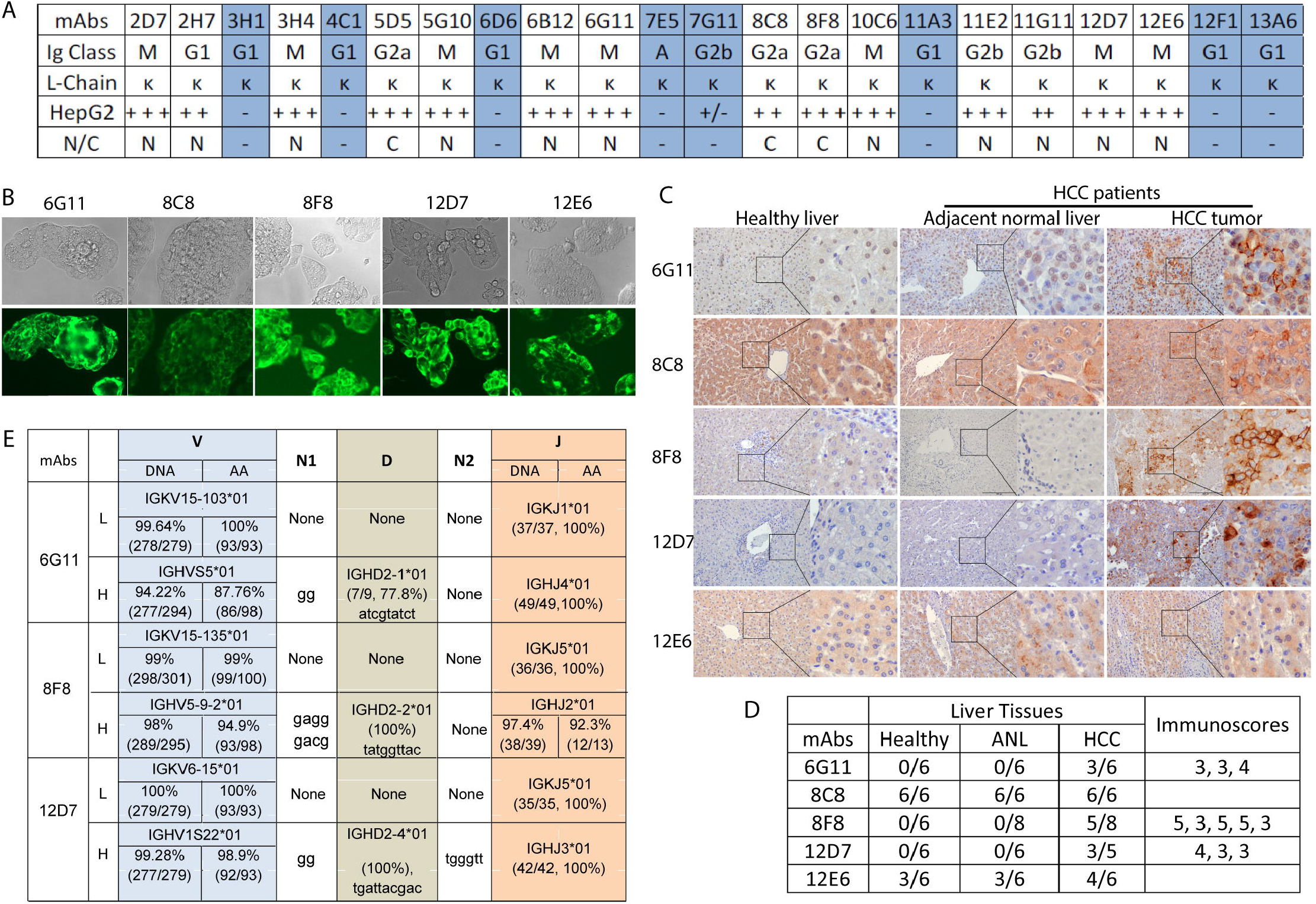
Generation and characterization of novel hGPC3-specifīc mAbs. (A) Summary of 22 new hGPC3 mAbs identified by ELISA assay with hGPC3 protein. The Ig class, staining of HepG2 cells, and binding of the hGPC3 N- or C-fragment were identified. The mAbs in blue shade did not stain HepG2 cells. (B) Representative photos of IF stainings of HepG2 cells by new hGPC3 mAbs. (C) Representative IHC staining of the paired HCC and the adjacent normal tissues of the same livers. Liver tissue from healthy donor was also included as control. A low and high magnifications were presented. Brown spots indicated positive staining. (D) Summary of IHC staining of multiple tissue samples by 5 mAbs. ANL: adjacent normal liver tissues. The immunoscores were calculated as described in “Supplementary Materials and Methods” and each number indicated one positive staining. The highest immunoscore is 8, the lowest score is 0. (E)The corresponding V, D, and J genes of the 6G11,8F8, and 12D7 cDNA were presented. Nucleotides added between V and D (N1) or D and J (N2) during recombination in the VH were also shown. The percentage and number in the parentheses indicate the percent identity and the length of nucleotides of indicated genes.

To further characterize the specificity, we purified 5 mAbs (6G11, 8C8, 8F8, 12D7, and 12E6) and conducted IHC staining of HCC and paired adjacent normal tissues. Healthy liver tissues were also included. We found that 3 mAbs (6G11, 8F8, and 12D7) specifically stained the HCC tissues but not the adjacent normal and healthy liver (Fig.1C-D and fig.S4). However, the other two mAbs had high non-specific staining of normal liver tissues. Thus, only 6G11, 8F8, and 12D7 were used in the following studies.

Next, the cDNA sequences of the heavy chain variable region (VH) and the light chain variable region (VL) of 6G11, 8F8, and 12D7 were identified and compared to the immunoglobulin (Ig) in the international ImMunoGeneTics (IMGT) databank (34). The results were summarized in Fig.1E.

### 6G11, 8F8, and 12D7 mAbs bind to different hGPC3 epitopes with distinct affinities

To search for low-affinity mAbs, we determined the binding affinity and epitopes of 6G11, 8F8, and 12D7, and compared them to GC33 (35). As the 4 mAbs belong to different Ig classes (6G11 and 12D7 are IgM, 8F8 is IgG2a, GC33 is IgG1), recombinant mAbs using VL and VH and a same human IgG H and L constant region were generated for fair comparison (Fig.2A). The recombinant mAbs were produced and purified from 293 cells (Fig.2B). The epitopes were mapped by Western blot and ELISA with overlapping peptides (fig.S5). The data showed that 6G11 and 8F8 bound to AA_25-39_ and AA_463-496_, respectively. The epitope of 8F8 is next to that of GC33 (30AA apart). However, no specific epitope was mapped for 12D7 although it bound to hGPC3-N fragment. 12D7 may recognizes a conformational epitope. The epitopes recognized by the mAbs were illustrated on the hGPC3 protein diagram (Fig.2C) (36). Next, the affinity of mAbs is determined by BioLayer Interferometry. The K_D_ of mAbs binding to hGPC3 were measured as 7.4nM for 6G11, 9.8nM for 12D7, 22.9nM for 8F8, and 1.38nM for GC33 (Fig.2D). The K_D_ of 8F8 is ~17 times higher than that of GC33. The *k*_on_ (association rate constant) of 8F8 and GC33 mAb binding hGPC3 are 8.97×10^3^/M^-1^S^-1^ and 5.56×10^4^M^-1^S^-1^, respectively, indicating that 8F8 mAb binds to hGPC3 is 6.2 times slower than GC33 mAb. The *k*_dis_ (dissociation rate constant or *k*_off_) of 8F8 and GC33 are 2.06×10^-4^/S^-1^ and 7.66×10^-5^/S^-1^, respectively, suggesting that 8F8-hGPC3 complex dissociate 2.7x times faster than GC33-hGPC3. Overall, compared to GC33, 8F8 mAb has a slow and unstable (Slow-on and Fast-off) engagement with hGPC3.

**Fig. 2.**
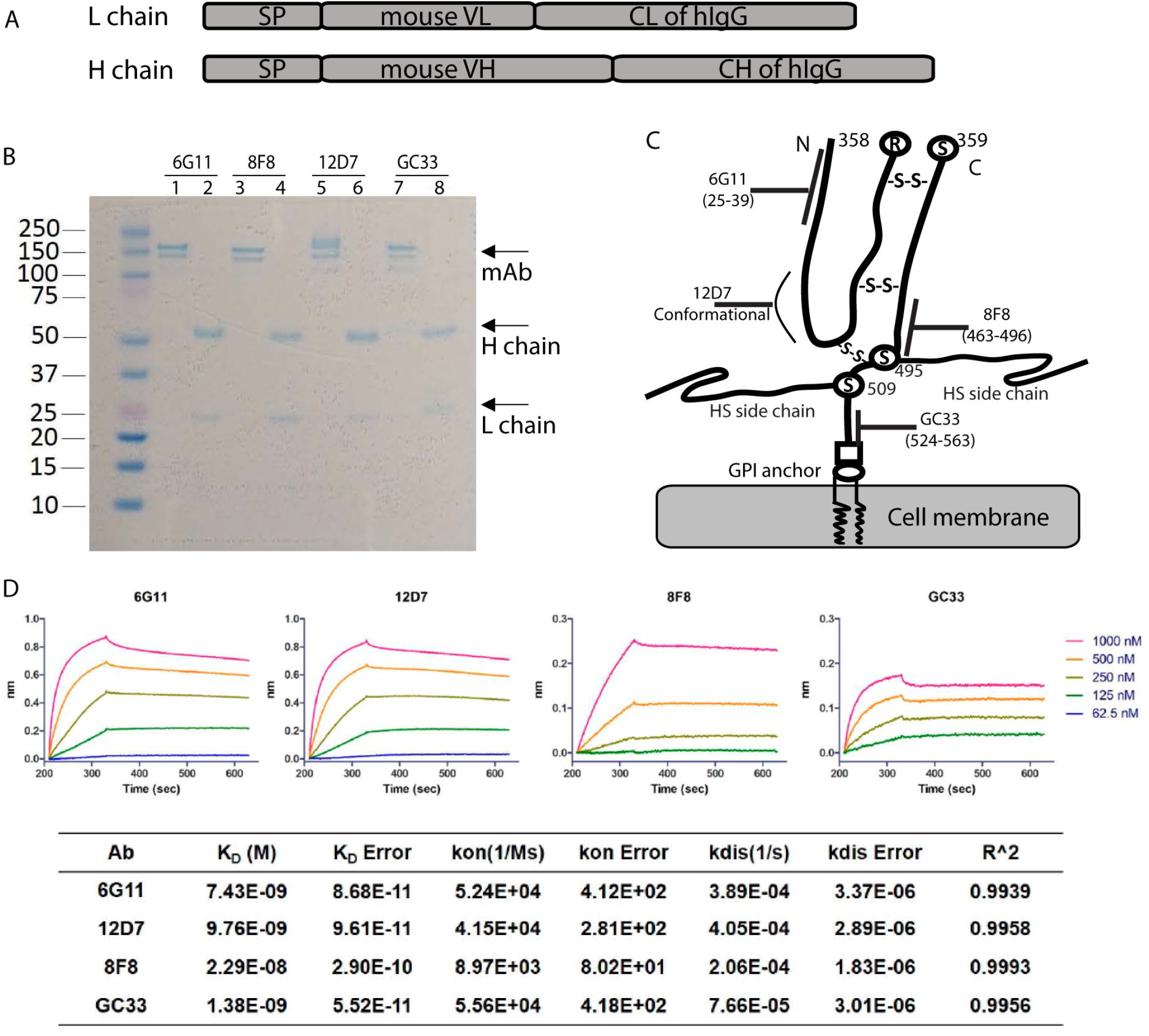
Identification of the binding epitopes and affinity of new hGPC3-specific mAbs. (A) The schematic structure of the recombinant mAbs. SP: signal peptide of human IgG;VL: variable region of light chain; VH: variable region of heavy chain; CL: constant region of human IgG light chain; CH: constant region of human IgG heavy chain. (B) PAGE analysis of the purified recombinant mAbs under reduced and non-reduced condition. mAbs were purified from the 293T cells after co-transfection of the L and H chain expressing plasmids. Gel was stained with SimplyBlue SafeStain. (C) The binding epitopes of mAbs are illustrated on a schematic structure of hGPC3 derived from Haruyama and Kataoka (ref 36). As a reference, GC33 mAb’s epitope was indicated. The potential disulfide bonds between N-fragment and C-fragment of hGPC3 were shown. (D) The K_D_, K_on_, and K_off_ value of each mAbs were determined by Biolayer interferometry. The measurement of affinity was repeated 3 times with similar data.

### 8F8 CART generates stronger antitumor effects than 6G11 and 12D7 CARTs

We hypothesized that 8F8 CART would generate stronger effects than 6G11 and 12D7 as reports showed that CARTs targeting membrane-proximal epitopes are more effective (37, 38). We built CARs using the single-chain variable fragment (scFv) of 6G11, 8F8, and 12D7 and 4-1BB co-stimulatory domain (Fig.3A). Human T cells were transduced by lentivectors to generate CARTs for *in vitro* and *in vivo* studies. CARs were approximately15-25% by anti-Fab staining (Fig.3B). All 3 CARTs could be expanded by HepG2 cells (fig.S6A), but only 6G11 and 8F8 CARTs killed HepG2 cells and produced IFNγ and IL-2 (Fig.3C). 6G11 and 8F8 CARTs were also able to kill hGPC3^lo^ Huh7 tumor cells (fig.S6B). 12D7 CART had no detectable cytotoxicity and cytokine. Secondly, compared to 6G11, 8F8 CART had stronger killing activity and produced more IL2 and IFNγ (Fig.3C).

**Fig.3.**
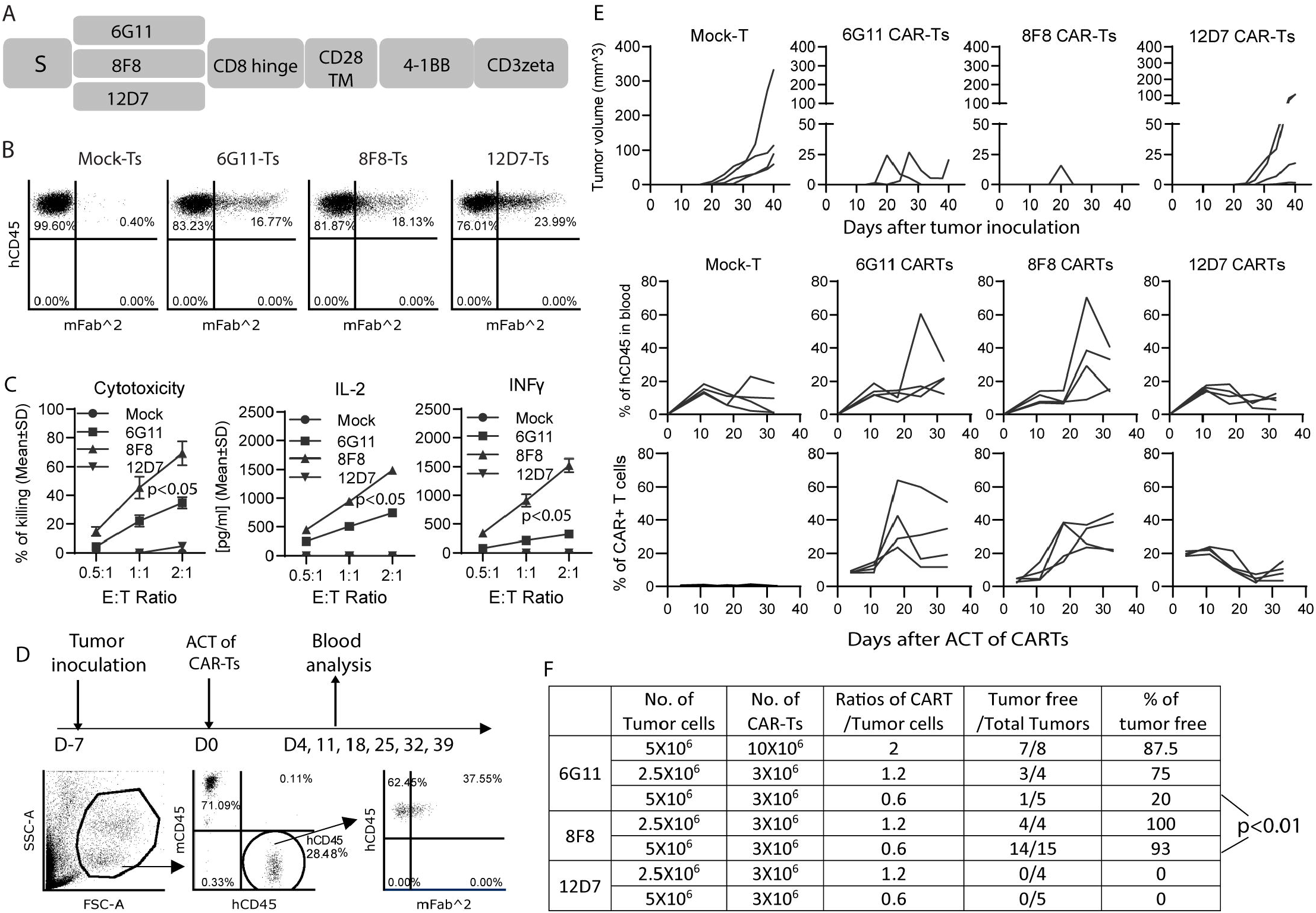
8F8 CARTs generate stronger antitumor effects than 6G11 and 12D7 CARTs. (A) Schematic structure of the 2nd generation CARs built from 6G11, 8F8, and 12D7. S: CD8 signal peptide, TM: CD28 transmembrane domain. Human T cells were isolated from donor PBMC and activated by CD3/CD28 beads on D −1. T cells were transduced on DO and beads were depleted on D1. CAR was detected on D7 by anti-Fab staining and in vitro assay was done between D7-10. (B) Representative dotplots of CARTs. (C) Cytotoxicity and cytokine production of CARTs after co-culture with HepG2 cells. Mean+/-SD was shown. Statistics were done with ANOVA. Experiment was repeated three times with similar observations. (D) The in vivo experiment plan and gating strategy of analyzing CARTs in the blood. Mouse endogenous mCD45 was stained as internal reference. (E) Individual tumor growth curve, the % of hCD45+, and % of CARTs after ACT. 4 or 5 mice were in each group. (F) Summary of the antitumor results from 3 experiments of treating HCC xenografts with different dose of CARTs. The numbers of tumor cells and CARTs were the initial numbers when they were injected into mice. The statistical analysis was done by NCSS software (Utah, USA).

To study antitumor effects, subcutaneous (SC) HCC xenografts in NSG mice were treated with CARTs (Fig.3D). 6G11 and 8F8 CARTs generated potent antitumor effects (Fig.3E and fig.S7). Consistent with *in vitro* data, no obvious antitumor effects was found in 12D7 CART group. The antitumor effects of 6G11 and 8F8 CARTs were dose-dependent (Fig.3F). Adoptive cell transfer (ACT) of 1.2-2 fold of 6G11 CART (compared to the number of inoculated HepG2 cells) resulted in complete tumor eradication in 75-90% of treated mice; but transfer of 0.6 fold of 6G11 CART generated only 20% of tumor free mice. In contrast, 8F8 CART resulted in complete eradication in more than 90% of the mice when only 0.6 fold of 8F8 CARTs was transferred. Correlating to its potent antitumor effects, 8F8 CART expanded more in the treated mice (Fig.3E). The group treated with 12D7 CARTs showed no increase of human T cells and CARTs. In summary, our data show that 8F8 CART targeting the membrane-proximal epitope generated the most potent antitumor effects, correlating to greater extent of *in vivo* expansion.

### 8F8 CARTs mediate similar *in vitro* effector functions as GC33 CARTs despite low avidity

As 8F8 and GC33 mAbs bind epitopes close to each other with 17 folds different affinity, in the following studies, we did comparative studies between 8F8 and GC33 CARTs. First, we conducted *in vitro* study. Because different affinity scFv may require different ICD for optimal activation (39), we constructed the CARs with CD28-CD3ζ (28z) or 4-1BB-CD3ζ (BBz) (Fig 4A). As anti-Fab staining did not clearly separate CARTs (Fig.3B), we tagged CARs with mCherry to reliably detect them and compare their level and hGPC3 binding avidity. Indeed, mCherry tag could clearly identify the CARTs even at very early time point of Day 3 after transduction (Fig.4B). The mCherry also allow better anti-Fab staining to separate the CART population. Multiple donor T cells were comparably transduced (fig.S8). Importantly, mCherry tag did not affect CART’s effector function (Fig.4C). Comparative study showed that, in both 28z and BBz ICD settings, 8F8 and GC33 CARTs had similar potent cytotoxicity and cytokine production (Fig.4D). The *in vitro* cytotoxicity was also assayed by Real-Time Cell Analyzer. Similar result was observed (fig.S9).

**Fig.4.**
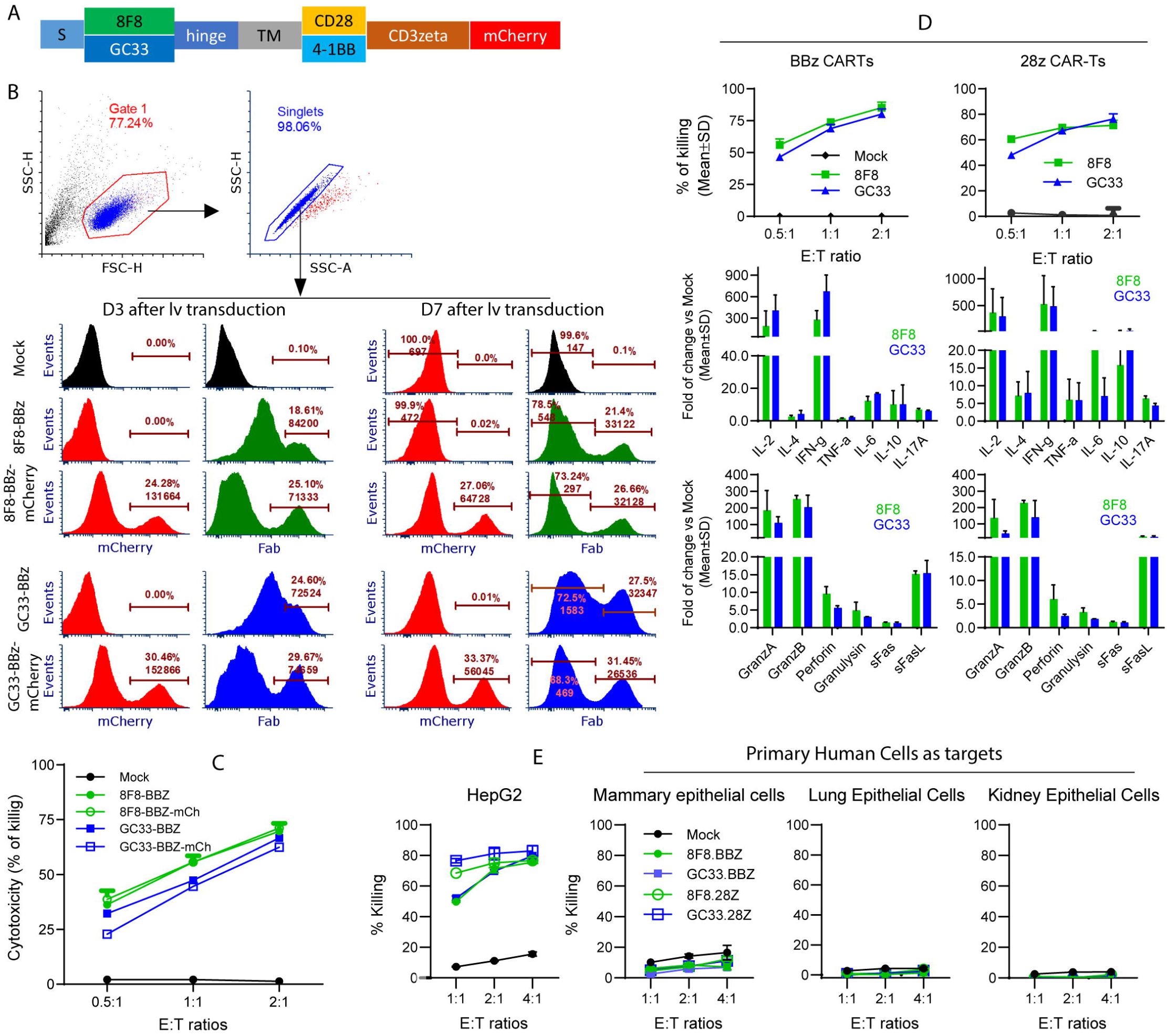
8F8 CARTs have similar in vitro effector function as GC33 CARTs despite low-avidity. (A) Schematic illustration of CARs with mCherry tag.The hinge is from CD8 andTM is from CD28. (B) mCherry tag enable clear identification of CAR+ cells. On D3 and D7 after transduction, cells were stained with anti-Fab antibody. Representative histogtam of BBz CARs were shown. (C) Comparison of the cytotoxicity of CARs with and without mCherry tag. Triplicate wells were used at each time point. The experiment was repeated 3 times with similar observation. (D) Cytotoxicity and cytokine production of different CARTs. The fold of increase over mock-T cells were presented. Each data was from the combined media of triplicate wells. Two Luminex testing repeats were conducted. (E) 8F8 and GC33 CARTs do not kill normal primary human cells. Primary human cells were co-cultured with CARTs for 24hrs and the media were used for LDH assay. HepG2 cells were used as positive control.

Next, to examine the CART’s off-tumor toxicity, we studied whether the 8F8 and GC33 CARTs could kill primary human mammary, lung, and kidney epithelial cells as they may express low level of hGPC3 (http://www.proteinatlas.org). The data showed that both 8F8 and GC33 CARTs did not kill those primary human cells (Fig.4E).

The mCherry tag enabled us to compare CART’s antigen binding avidity. First, by mCherry intensity, we showed similar level of 8F8 and GC33 CARs although 28z CAR was slightly (1.3 times) higher than BBz CAR (fig.S10). Secondly, 8F8 CARTs with BBz or 28z ICD bound 4-5 times less of hGPC3 protein than counterpart GC33 CARTs, suggesting 8F8 CAR is indeed low-avidity. Thirdly, 28z CARTs bound more than 2 times of hGPC3 than BBz CART even though their CAR level was only slightly (1.3 times) higher. Our data indicate that the 8F8 CAR is indeed low-avidity compared to GC33 CAR and that the BBz CAR binds less hGPC3 than 28z CAR.

### 8F8-BBz CART mediates durable antitumor effect with enhanced *in vivo* expansion

To study whether low-avidity CART is superior to high avidity CARTs *in vivo*, we utilized the intraperitoneal (IP) HCC xenografts to study the *in vivo* expansion and antitumor effects of 8F8 and GC33 CARTs with BBz or 28z ICD. HepG2 cells expressing luciferase (HepG2-Luc) were generated. Three million of HepG2-Luc cells were inoculated into the peritoneal cavity to establish IP tumors for 10 days before treatment (Fig. 5A). Two weeks after ACT, nearly every tumor in all CART treated groups was eliminated (Fig.5B-5C). However, 40% of the mice in the GC33-28z, 8F8-28z, and GC33-BBz treated groups showed tumor regrowth 4-5 weeks after ACT. In contrast, no tumor re-growth was observed in 8F8-BBz group. In addition, significantly more human T cells and CARTs were found in the 8F8 CART treated mice, especially in the 8F8-BBz CART group (Fig.5D).

**Fig. 5.**
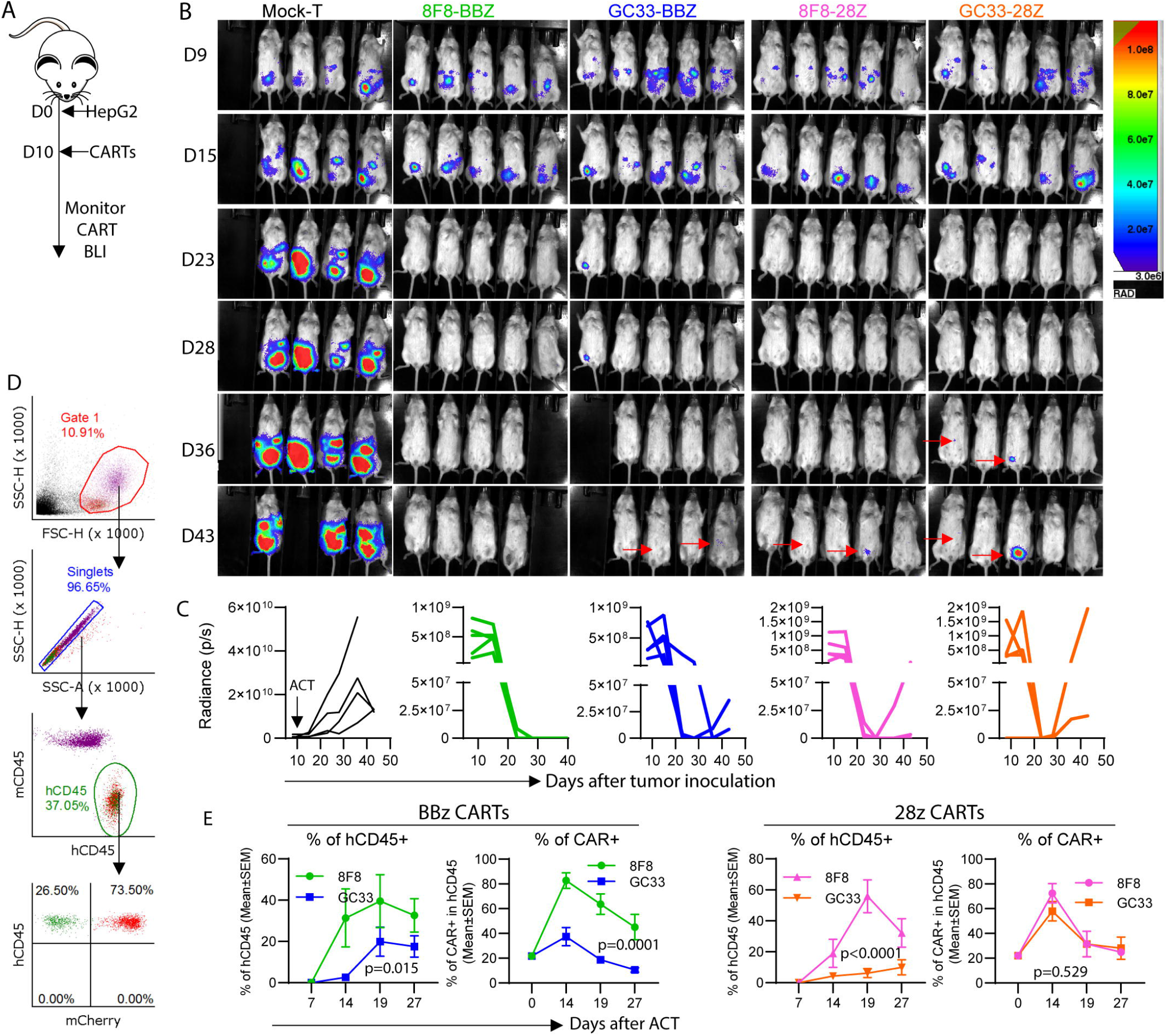
8F8-BBz CART generates durable antitumor effect with enhanced expansion and persistence. (A) Experimental plan. (B) Bioluminescent imaging (BLI) of the IP tumors. All images at different time points and from different groups were adjusted to the same scale. (C) Tumor BLI kinetics of each individual mouse, 5 mice in each treated group. (D and E) Gating strategy and detection of hCD45 T cells and CAR+ T cells in the blood. Endogenous mCD45÷ cells were used as reference. Statistical analysis was done with SAS System GLM procedure.

To compare the 8F8 and GC33 CARTs further, we treated larger (22-day) IP tumors and investigated CART’s *in vivo* expansion (fig.S11A). Although CARTs were unable to eradicate such large tumors (fig.S11B), the survival of the mice treated with 8F8-BBz CART was significantly longer than other groups (fig.S11C). Again, the % of hCD45+ T cells and % of CARTs were significantly higher and persisted longer in 8F8-BBz CART group than other groups (fig.S11D). This data suggests that, compared to GC33 and 8F8-28z CARTs, 8F8-BBz CART generates enhanced expansion and persistence, especially in the presence of large tumor burden, resulting in durable antitumor effects and extending survival.

### More 8F8-BBz CARTs infiltrate and persist in the tumor mass than GC33 CARTs

We next examined tumor-infiltrating CARTs to test the hypothesis that low-avidity 8F8 CART expand more and persists longer in solid tumors. To obtain sufficient tumor-infiltrating CARTs, we established large SC tumors (4 weeks, ~1cm in diameter) before ACT (Fig.6A). First, the hCD45 cells in blood were similar (30-50%) on Day 12 after ACT in all CART groups. On Day 16, the hCD45+ cells in 8F8-BBz and GC33-BBz CART groups increased 3-folds (30% to 90%) and 2-folds (from 40% to 80%), respectively (Fig.6B). However, while the hCD45+ T cells in the 8F8-28z CART group modestly increased (30% to 48%), it slightly decreased in the GC33-28z group (45% to 42%). Secondly, BLI measurement showed 8F8-BBz CART generated more durable tumor control compared to other CARTs (Fig.6C), similar to the antitumor effects observed in smaller IP tumors (Fig.5). Remarkably, there were far more tumor-infiltrating 8F8-BBz CARTs than any other CARTs, especially in later time points (Fig.6D). Nearly all live cells in the single cell suspension of the BBz CART treated tumor were CAR+ (Fig.6E). On the other hand, the single cell suspension of the GC33-28z treated group contained more non-hCD45 T cells (likely HepG2 tumor cells). The CART enumeration in the tumor lesion showed that there were 5 times more of 8F8-BBz CARTs than other CART groups (Fig.6F).

**Fig.6.**
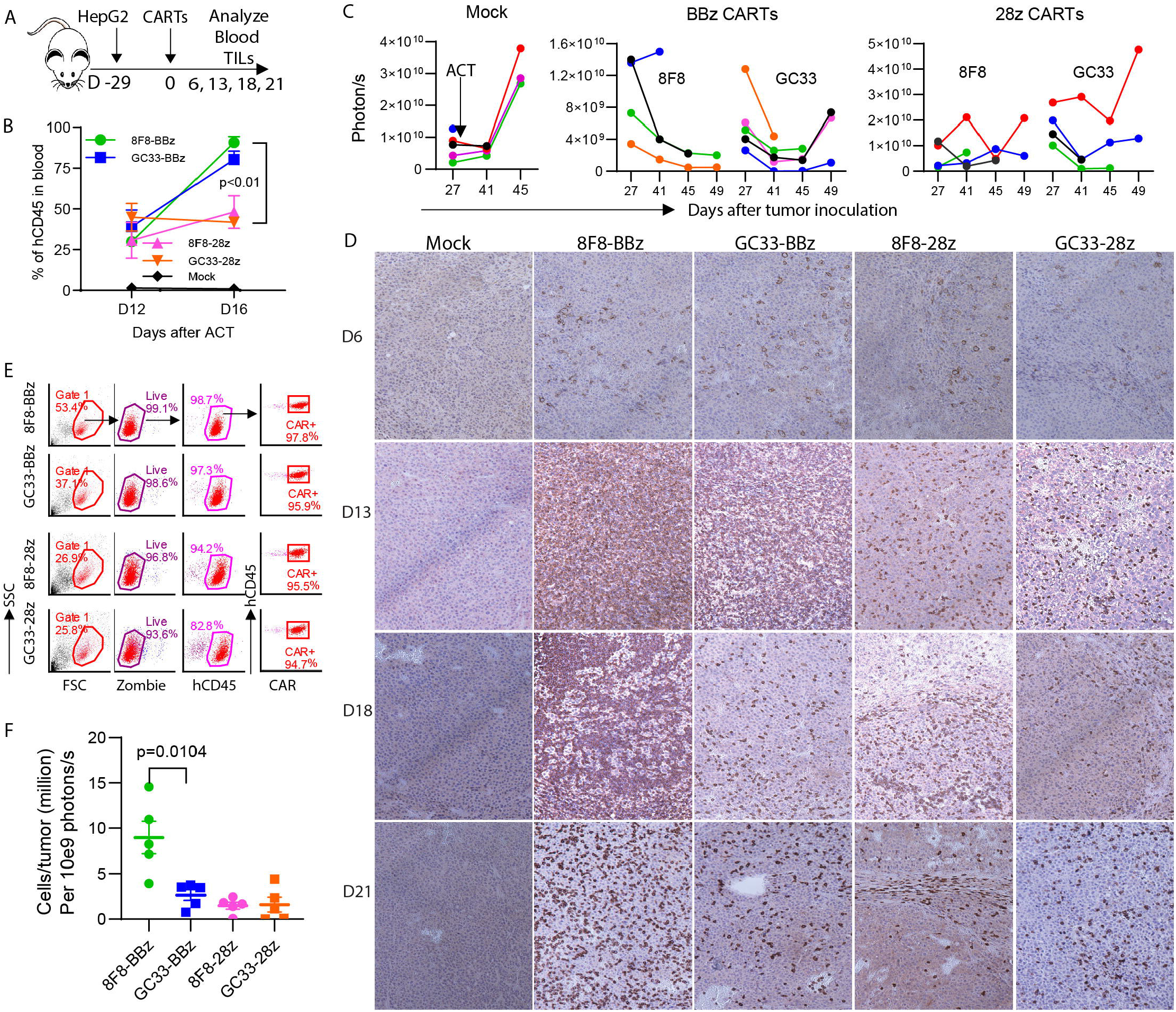
8F8-BBz CARTs infiltrate more and persist longer in the tumor mass than other CARTs. (A) Experimental plan. (B) The change of hCD45+T cells in the blood from DI 2 to DI 6 after ACT. (C) Tumor BLI kinetics after CART treatment. Each line represented an individual tumor. (D) Representative IHC staining of tumor infiltraing CD8T cells at indicated time points after ACT. To cover more area, small magnification lense (1 OX) was used. (E) The gating strategy for enumerating the tumor infiltrating CARTs. The viable cells from the single tumor cell suspension were counted by Trypan blue. The CART number in the tumor was calculated by the total viable cells in the tumor x% of hCD45 x %of CAR+. (F) Summary of the tumor infiltrating CART numbers of 5 mice from 3 time points (DI 3, D18 and D2l).The number was normalized by dividing the total CART number in the tumor by the corresponding tumor mass BLI. The experiment was repeated twice and similar data was observed.

### 8F8-BBz CARTs resist exhaustion and maintain better function in tumors

In this experiment, we tested the hypothesis that low-avidity CARTs were more resistant to exhaustion and apoptosis and maintained better effector function in solid tumors. To this end, we studied the exhaustion, apoptosis, and effector function of the tumor-infiltrating CARTs. The PD1, LAG3, and Annexin-V staining showed that BBz CARTs were less exhausted and apoptotic than 28z CARTs and that low-avidity 8F8 CARTs were less exhausted and apoptotic than high-avidity GC33 CARTs (Fig.7A-B and fig.S12A). The 8F8-BBz and GC33-28z CART were the least and the most exhausted and apoptotic CARTs, respectively (Fig.7B). The loss of cytotoxicity function in the tumor was CART and time dependent (Fig.7C and fig.S12B). In agreement with their high exhaustion and apoptosis, 28z CARTs in tumor lost their killing activity faster than BBz CARTs. The low-affinity 8F8 CARTs retained effector function longer than GC33 CARTs. The tumor-infiltrating 8F8-BBz CART maintained substantial cytotoxicity at time point of Day 21 after treatment when other CART’s function were completely lost or severely compromised (Fig.7C). In addition, the tumor-infiltrating CARTs were sorted and their gene expression profile was analyzed by bulk RNAseq. The gene profile data was in agreement with immunological data. Differential gene expression between 8F8 and GC33 CARTs were observed (Fig.7D and fig.S12C). The genes related to T cell exhaustion is lower in 8F8 than in GC33 CARTs. In contrast, expression of genes related to effector function, memory, and stem-like memory were higher in 8F8 CARTs than in GC33 CARTs. In conclusion, more 8F8-BBz CARTs are found in the solid tumor lesions, they are less exhausted and apoptotic, and maintain better effector function than 8F8-28z and high-avidity GC33 CARTs.

**Fig.7.**
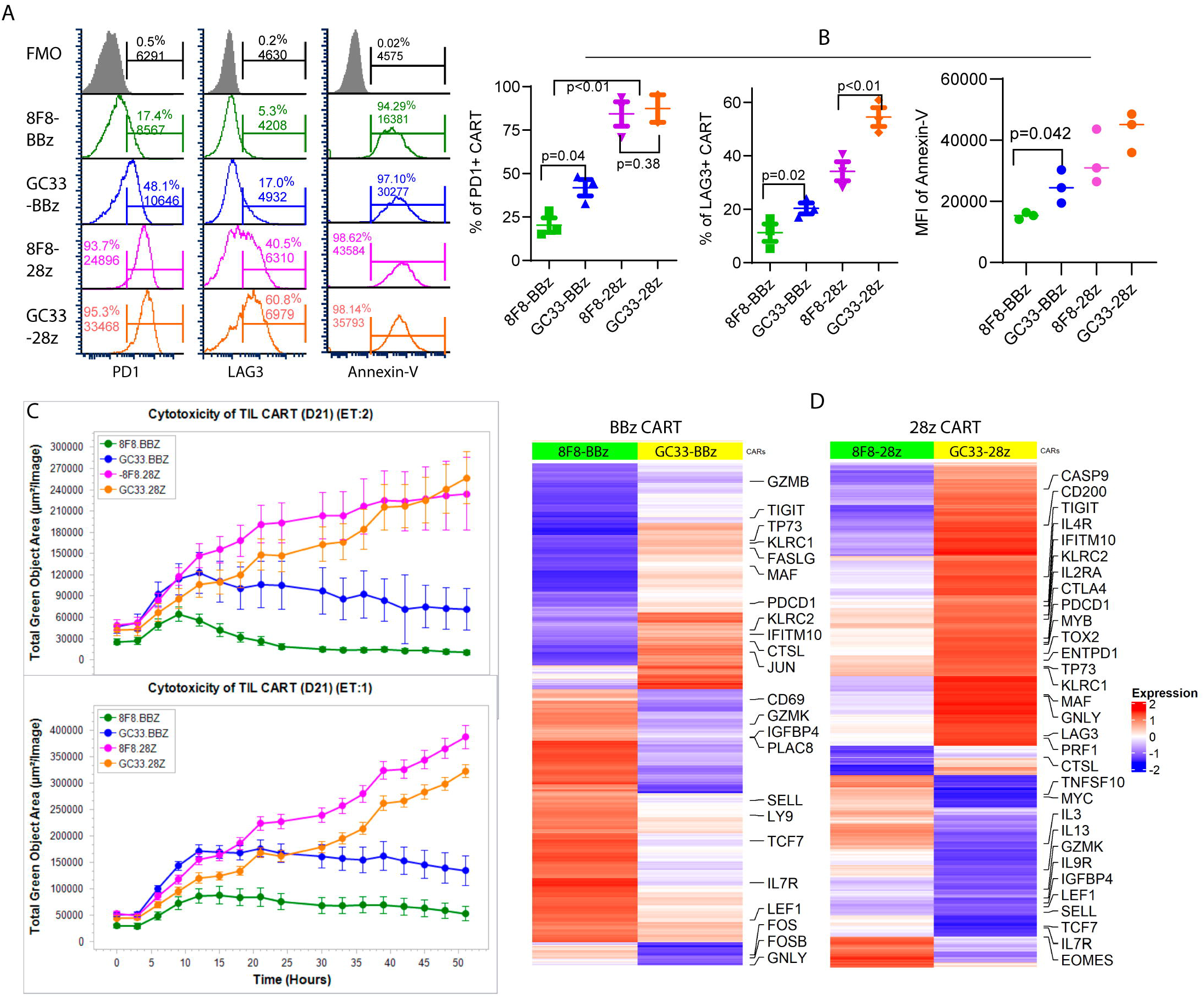
8F8-BBz CARTs in tumors are less exhausted and more functional. (A) The exhaustion and apoptosis of tumor-infiltrating CARTs at D21 after ACT. The tumor infiltrating CARTs from the single cell suspensions were stained and gated as Fig 6E.The CARTs were gated and analyzed for PD1, LAG3, or Annexin-V. FMO (Fluorescence Minus One) was used to set up the gate. (B) Summary of the exhaustion and apoptosis of tumor infiltrating CARTs. The % of PD1 + and LAG+ of CARTs, and the MFI of Annexin V on CARTs were summarized from 3 mice. (C) Cytotoxicity of CARTs isolated from tumors on D21 after ACT. Each data point was calculated from 10 datasets of the HepG2-GFP area from 2 wells. The experiment was repeated twice with similar observation. (D) Heatmaps show differentially expressed genes between 8F8 and GC33 CART cells. Genes coordinately up or down regulated at least 1,2-fold between 8F8 and GC33 CART cells were plotted in the heatmaps. Labels highlight genes involved in memory and effector differentiation or exhaustion, and apoptosis pathways.

### 8F8 CARTs are less exhausted and apoptotic after *in vitro* co-culture with tumor cells

We then conducted in vitro co-culture experiments to further study whether low-avidity 8F8 CART is less exhausted than GC33 CART after engaging with target tumor cells (Fig.8A). **1)** Our data showed that the % of CART decreased during their initial 1-3 days co-culture with HepG2 cells (Fig.8B). The decrease of 8F8 CART was mild compared to GC33 CART. This observation is in agreement with previous report that CART could be depleted by tumor cells (29). After 2-3 days, the % of CARTs began to recover and the 8F8 CART came back faster than GC33 CART. **2)** Compared to GC33, the 8F8 CART was much less apoptotic after interacting with HepG2 cells (Fig.8C). The MFI of Annexin V on 8F8-BBz CART was 5 times lower than GC33-BBz CART. **3)** The 8F8 CART also expressed less PD1 and LAG3. Both the % and MFI of PD1 and LAG3 were lower on 8F8 CART than on GC33 CART (Fig.8D). The LAG3 and PD1 double positive cells were less in 8F8 CART than GC33 CART (Fig.8E). **4)** The 8F8 CART contained more CCR7 and CD62L double positive central memory cells and CD45RA and CD62L double positive naïve-like T cells after interacting with tumor cells (Fig.8F). The same pattern of less apoptosis and exhaustion in 8F8-BBz CART was also observed in 8F8-28z CART than GC33-28z CART (fig.S13). The CART’s exhaustion and apoptosis were more severe when more tumor cells are present (fig.S13), further suggesting that the CART apoptosis and exhaustion are driven by antigen engagement and stimulation. In summary, the *in vitro* study supports *in vivo* findings and suggests the low-avidity CARTs are less prone to apoptosis and exhaustion and can preserve memory-like property, thus will persist in the presence of constant antigen stimulation in solid tumors to achieve stronger and durable antitumor effect.

**Fig.8.**
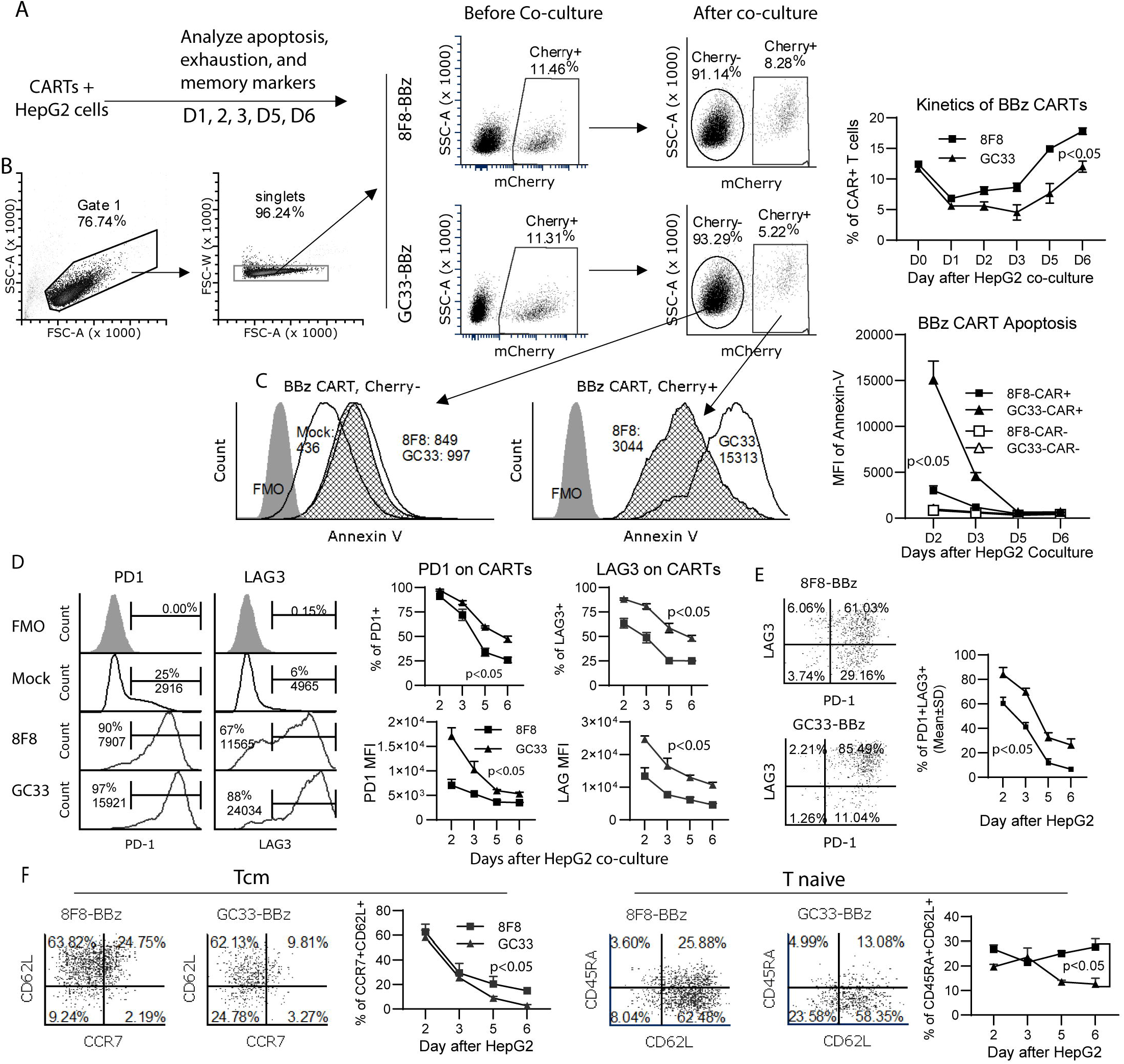
8F8 CARTs are less exhausted and apoptotic, and contain more memory cells after co-culture with tumor cells. (A) THe experimental scheme. (B) The gating strategy and kinetics of CAR+ changes were shown. (C) The apoptosis staining of CAR- and CAR+T cells after HepG2 stimulation.The numbers in the figure indicate the MFI of Anne×in-VThe dynamic changes of apoptosis was also summarized. (D and E)The %and MFI of exhaustion markers, and the %of PD1 and LAG3 double positive cells among the CARTs after HepG2 stimulation were presented. (F). The % of central memory (Tern) and naive-like cells (Tnaive) cells in the CAR+ cells after HepG2 co-culture were shown. Each data point represents mean+/-SD of triplicate wells. The experiments were repeated 3 times with similar observation.

## Discussion

In this study, we developed a low-affinity 8F8 mAb and created low-avidity CARTs. We found that, compared to high-avidity GC33 CARTs, the low-avidity 8F8-BBz CART generated enhanced expansion and persistence *in vivo*. More 8F8-BBz CARTs infiltrated into tumor lesions, resulting in potent and durable antitumor effects, which correlates to its resistance to exhaustion and maintenance of effector function in tumor lesions. As loss of effector function is one of the main obstacles to the success of immunotherapy, low-avidity CARTs with resistance to exhaustion may benefit a wide range of solid tumors. The findings made in this study may encourage more investigations to develop low-avidity CARTs that eventually lead to success of solid tumor therapy.

The engagement avidity and duration likely determine the fate of both CARTs and tumor cells. The engagement avidity needs to reach certain level to kill tumor cells, but above a threshold does not increase CART efficacy (40), but instead may drive CART to exhaustion. The low-avidity CART with faster dissociation rate *k*_off_ will easily disengage from tumor cells. Such “fly-kiss” style engagement may give CART a short break to avoid exhaustion and preserve its function inside tumor lesion. This is in consistent with recent reports that controlled CAR expression by synthetic Notch circuit (41, 42) only after engaging with tumor cells and transient inducible shut-off of CAR expression with drugs (43) enhanced CART’s persistence and antitumor effects. In both cases, CARTs have a break time to avoid exhaustion.

ICD domains also affect CART activation (39). In addition to stronger signaling, we found that the 28z CAR bound 2 times more of antigen than BBz CAR. The stronger avidity and signaling of CD28z results in more severe exhaustion, apoptosis, and loss of effector function of 28z CART than BBz CART. Among the 4 CARTs, GC33-28z CART has the highest avidity and the highest exhaustion and apoptosis, and has the fastest loss of effector function. This data may explain why the antitumor efficacy of GC33-28z CART in clinical trial is low (25). As the CAR signaling strength above certain threshold may not be necessary but rather detrimental (44), CARs should be optimized by finely balancing the desired signaling strength, anti-tumor response, and potential risk of off-tumor toxicity. Lastly, affinity thresholds are likely not universal and depend on interconnected factors such as antigen densities on target cells, CAR expression levels, and binding epitope location (45, 46). Such efforts may inspire others to develop CARTs that have stronger resistance to antigen-driven exhaustion and apoptosis and maintain better effector function to treat solid tumors.

High-avidity CARTs may generate a rapid antitumor response, but they are often shortlived in tumor lesions, resulting in transient antitumor effects. Unless CARTs completely eradicate tumors before they are exhausted, tumors will relapse. On the other hand, the antitumor effects of low-avidity CARTs is slow, but their long survival and maintenance of effector function in the tumor lesion ensure them to have durable antitumor effects. However, we also found that, although the low-avidity 8F8-BBz CART significantly extended mouse survival, it did not eradicate large tumors (>22 day IP tumors). Thus, in addition to exhaustion-resistant low-avidity CARTs, modulation of tumor microenvironment may be also critical to the success of solid cancer immunotherapy (47). The level of PD1 and LAG3 on tumor-infiltrating 8F8 CART is 2-3 fold lower than GC33 CARTs. The low-avidity CARTs stay in a “lightly exhausted” state (PD^lo^), and be prevented to become “deeply exhausted” (PD^hi^) (48) in the tumor. Previous reports showed that the effector function of this PD^lo^, but not PD^hi^ T cells is reversible by checkpoint blockade (49). Thus, it will be interesting to study if combination of checkpoint blockade will further enhance the antitumor effects of low-avidity CARTs. GPC3 is detected in ~70% of HCC and 50% of squamous lung cancers, but is not detected in healthy tissues (50), the low-avidity 8F8-BBz CART should have a broad application and will likely generate better therapeutic effects in treating HCC and other hGPC3+ solid tumors.

## References

1. Sung H, Ferlay J, Siegel RL, Laversanne M, Soerjomataram I, Jemal A, Bray F. Global cancer statistics 2020: GLOBOCAN estimates of incidence and mortality worldwide for 36 cancers in 185 countries. CA Cancer J Clin 2021.

2. Pico de Coana Y, Choudhury A, Kiessling R. Checkpoint blockade for cancer therapy: revitalizing a suppressed immune system. Trends Mol Med 2015;21:482–491.

3. Zhu W, Peng Y, Wang L, Hong Y, Jiang X, Li Q, Liu H, et al. Identification of alpha-fetoprotein-specific T-cell receptors for hepatocellular carcinoma immunotherapy. Hepatology 2018;68:574–589.

4. Docta RY, Ferronha T, Sanderson JP, Weissensteiner T, Pope GR, Bennett AD, Pumphrey NJ, et al. Tuning T-Cell Receptor Affinity to Optimize Clinical Risk-Benefit When Targeting Alpha-Fetoprotein-Positive Liver Cancer. Hepatology 2018.

5. June CH, Sadelain M. Chimeric Antigen Receptor Therapy. N Engl J Med 2018;379:64–73.

6. Caraballo Galva LD, Cai L, Shao Y, He Y. Engineering T cells for immunotherapy of primary human hepatocellular carcinoma. J Genet Genomics 2020;47:1–15.

7. Wherry EJ. T cell exhaustion. Nat Immunol 2011;12:492–499.

8. Thommen DS, Schumacher TN. T Cell Dysfunction in Cancer. Cancer Cell 2018;33:547–562.

9. Hombach A, Hombach AA, Abken H. Adoptive immunotherapy with genetically engineered T cells: modification of the IgG1 Fc ‘spacer’ domain in the extracellular moiety of chimeric antigen receptors avoids ‘off-target’ activation and unintended initiation of an innate immune response. Gene Ther 2010;17:1206–1213.

10. Long AH, Haso WM, Shern JF, Wanhainen KM, Murgai M, Ingaramo M, Smith JP, et al. 4-1BB costimulation ameliorates T cell exhaustion induced by tonic signaling of chimeric antigen receptors. Nature Medicine 2015;21:581–590.

11. Gargett T, Yu W, Dotti G, Yvon ES, Christo SN, Hayball JD, Lewis ID, et al. GD2-specific CAR T Cells Undergo Potent Activation and Deletion Following Antigen Encounter but can be Protected From Activation-induced Cell Death by PD-1 Blockade. Mol Ther 2016;24:1135–1149.

12. D’Aloia MM, Zizzari IG, Sacchetti B, Pierelli L, Alimandi M. CAR-T cells: the long and winding road to solid tumors. Cell Death Dis 2018;9:282.

13. Majzner RG, Mackall CL. Clinical lessons learned from the first leg of the CAR T cell journey. Nat Med 2019;25:1341–1355.

14. Jayaraman J, Mellody MP, Hou AJ, Desai RP, Fung AW, Pham AHT, Chen YY, et al. CAR-T design: Elements and their synergistic function. EBioMedicine 2020;58:102931.

15. Hudecek M, Lupo-Stanghellini MT, Kosasih PL, Sommermeyer D, Jensen MC, Rader C, Riddell SR. Receptor affinity and extracellular domain modifications affect tumor recognition by ROR1-specific chimeric antigen receptor T cells. Clin Cancer Res 2013;19:3153–3164.

16. Lynn RC, Feng Y, Schutsky K, Poussin M, Kalota A, Dimitrov DS, Powell Jr DJ. High-affinity FRß-specific CAR T cells eradicate AML and normal myeloid lineage without HSC toxicity. Leukemia 2016;30:1355–1364.

17. Richman SA, Nunez-Cruz S, Moghimi B, Li LZ, Gershenson ZT, Mourelatos Z, Barrett DM, et al. High-Affinity GD2-Specific CAR T Cells Induce Fatal Encephalitis in a Preclinical Neuroblastoma Model. Cancer Immunol Res 2018;6:36–46.

18. Ghorashian S, Kramer AM, Onuoha S, Wright G, Bartram J, Richardson R, Albon SJ, et al. Enhanced CAR T cell expansion and prolonged persistence in pediatric patients with ALL treated with a low-affinity CD19 CAR. Nat Med 2019;25:1408–1414.

19. Gao H, Li K, Tu H, Pan X, Jiang H, Shi B, Kong J, et al. Development of T cells redirected to glypican-3 for the treatment of hepatocellular carcinoma. Clin Cancer Res 2014;20:6418–6428.

20. Li D, Li N, Zhang YF, Fu H, Feng M, Schneider D, Su L, et al. Persistent Polyfunctional Chimeric Antigen Receptor T Cells That Target Glypican 3 Eliminate Orthotopic Hepatocellular Carcinomas in Mice. Gastroenterology 2020;158:2250–2265 e2220.

21. Liu X, Gao F, Jiang L, Jia M, Ao L, Lu M, Gou L, et al. 32A9, a novel human antibody for designing an immunotoxin and CAR-T cells against glypican-3 in hepatocellular carcinoma. J Transl Med 2020;18:295.

22. Baumhoer D, Tornillo L, Stadlmann S, Roncalli M, Diamantis EK, Terracciano LM. Glypican 3 expression in human nonneoplastic, preneoplastic, and neoplastic tissues: a tissue microarray analysis of 4,387 tissue samples. Am J Clin Pathol 2008;129:899–906.

23. Nakano K, Orita T, Nezu J, Yoshino T, Ohizumi I, Sugimoto M, Furugaki K, et al. Anti-glypican 3 antibodies cause ADCC against human hepatocellular carcinoma cells. Biochem Biophys Res Commun 2009;378:279–284.

24. Phung Y, Gao W, Man YG, Nagata S, Ho M. High-affinity monoclonal antibodies to cell surface tumor antigen glypican-3 generated through a combination of peptide immunization and flow cytometry screening. MAbs 2012;4:592–599.

25. Shi D, Shi Y, Kaseb AO, Qi X, Zhang Y, Chi J, Lu Q, et al. Chimeric Antigen Receptor-Glypican-3 T-Cell Therapy for Advanced Hepatocellular Carcinoma: Results of Phase I Trials. Clin Cancer Res 2020;26:3979–3989.

26. Liu X, Jiang S, Fang C, Yang S, Olalere D, Pequignot EC, Cogdill AP, et al. Affinity-Tuned ErbB2 or EGFR Chimeric Antigen Receptor T Cells Exhibit an Increased Therapeutic Index against Tumors in Mice. Cancer Res 2015;75:3596–3607.

27. Drent E, Themeli M, Poels R, de Jong-Korlaar R, Yuan H, de Bruijn J, Martens ACM, et al. A Rational Strategy for Reducing On-Target Off-Tumor Effects of CD38-Chimeric Antigen Receptors by Affinity Optimization. Mol Ther 2017;25:1946–1958.

28. Park S, Shevlin E, Vedvyas Y, Zaman M, Park S, Hsu YS, Min IM, et al. Micromolar affinity CAR T cells to ICAM-1 achieves rapid tumor elimination while avoiding systemic toxicity. Sci Rep 2017;7:14366.

29. Hoseini SS, Dobrenkov K, Pankov D, Xu XL, Cheung NK. Bispecific antibody does not induce T-cell death mediated by chimeric antigen receptor against disialoganglioside GD2. Oncoimmunology 2017;6:e1320625.

30. Xiao H, Zhu P, Liu B, Pan Q, Jiang X, Xu X, Fu N. Generation and characterization of human delta-globin-specific monoclonal antibodies. Blood Cells Mol Dis 2010;44:127–132.

31. Walchli S, Loset GA, Kumari S, Johansen JN, Yang W, Sandlie I, Olweus J. A practical approach to T-cell receptor cloning and expression. PLoS One 2011;6:e27930.

32. He Y, Zhang J, Donahue C, Falo LD, Jr. Skin-derived dendritic cells induce potent CD8(+) T cell immunity in recombinant lentivector-mediated genetic immunization. Immunity 2006;24:643–656.

33. He Y, Zhang J, Mi Z, Robbins P, Falo LD, Jr. Immunization with lentiviral vector-transduced dendritic cells induces strong and long-lasting T cell responses and therapeutic immunity. J Immunol 2005;174:3808–3817.

34. Lefranc MP, Giudicelli V, Duroux P, Jabado-Michaloud J, Folch G, Aouinti S, Carillon E, et al. IMGT(R), the international ImMunoGeneTics information system(R) 25 years on. Nucleic Acids Res 2015;43:D413–422.

35. Ishiguro T, Sugimoto M, Kinoshita Y, Miyazaki Y, Nakano K, Tsunoda H, Sugo I, et al. Anti-glypican 3 antibody as a potential antitumor agent for human liver cancer. Cancer Res 2008;68:9832–9838.

36. Haruyama Y, Kataoka H. Glypican-3 is a prognostic factor and an immunotherapeutic target in hepatocellular carcinoma. World J Gastroenterol 2016;22:275–283.

37. Hombach AA, Schildgen V, Heuser C, Finnern R, Gilham DE, Abken H. T cell activation by antibody-like immunoreceptors: the position of the binding epitope within the target molecule determines the efficiency of activation of redirected T cells. J Immunol 2007;178:4650–4657.

38. James SE, Greenberg PD, Jensen MC, Lin Y, Wang J, Till BG, Raubitschek AA, et al. Antigen sensitivity of CD22-specific chimeric TCR is modulated by target epitope distance from the cell membrane. J Immunol 2008;180:7028–7038.

39. Drent E, Poels R, Ruiter R, van de Donk N, Zweegman S, Yuan H, de Bruijn J, et al. Combined CD28 and 4-1BB Costimulation Potentiates Affinity-tuned Chimeric Antigen Receptor-engineered T Cells. Clin Cancer Res 2019;25:4014–4025.

40. Chmielewski M, Hombach A, Heuser C, Adams GP, Abken H. T cell activation by antibody-like immunoreceptors: increase in affinity of the single-chain fragment domain above threshold does not increase T cell activation against antigen-positive target cells but decreases selectivity. J Immunol 2004;173:7647–7653.

41. Choe JH, Watchmaker PB, Simic MS, Gilbert RD, Li AW, Krasnow NA, Downey KM, et al. SynNotch-CAR T cells overcome challenges of specificity, heterogeneity, and persistence in treating glioblastoma. Sci Transl Med 2021;13.

42. Hyrenius-Wittsten A, Su Y, Park M, Garcia JM, Alavi J, Perry N, Montgomery G, et al. SynNotch CAR circuits enhance solid tumor recognition and promote persistent antitumor activity in mouse models. Sci Transl Med 2021;13.

43. Weber EW, Parker KR, Sotillo E, Lynn RC, Anbunathan H, Lattin J, Good Z, et al. Transient rest restores functionality in exhausted CAR-T cells through epigenetic remodeling. Science 2021;372.

44. Künkele A, Johnson AJ, Rolczynski LS, Chang CA, Hoglund V, Kelly-Spratt KS, Jensen MC. Functional Tuning of CARs Reveals Signaling Threshold above Which CD8^+^CTL Antitumor Potency Is Attenuated due to Cell Fas–FasL-Dependent AICD. Cancer Immunology Research 2015;3:368–379.

45. Frigault MJ, Lee J, Basil MC, Carpenito C, Motohashi S, Scholler J, Kawalekar OU, et al. Identification of chimeric antigen receptors that mediate constitutive or inducible proliferation of T cells. Cancer Immunol Res 2015;3:356–367.

46. Arcangeli S, Rotiroti MC, Bardelli M, Simonelli L, Magnani CF, Biondi A, Biagi E, et al. Balance of Anti-CD123 Chimeric Antigen Receptor Binding Affinity and Density for the Targeting of Acute Myeloid Leukemia. Mol Ther 2017;25:1933–1945.

47. Fuca G, Reppel L, Landoni E, Savoldo B, Dotti G. Enhancing Chimeric Antigen Receptor T-Cell Efficacy in Solid Tumors. Clin Cancer Res 2020;26:2444–2451.

48. Xu-Monette ZY, Zhang M, Li J, Young KH. PD-1/PD-L1 Blockade: Have We Found the Key to Unleash the Antitumor Immune Response? Front Immunol 2017;8:1597.

49. Philip M, Fairchild L, Sun L, Horste EL, Camara S, Shakiba M, Scott AC, et al. Chromatin states define tumour-specific T cell dysfunction and reprogramming. Nature 2017;545:452–456.

50. Yamauchi N, Watanabe A, Hishinuma M, Ohashi K, Midorikawa Y, Morishita Y, Niki T, et al. The glypican 3 oncofetal protein is a promising diagnostic marker for hepatocellular carcinoma. Mod Pathol 2005;18:1591–1598.

