## Supplemental materials and data for "Novel low-avidity glypican-3 specific CARTs resist exhaustion and mediate durable antitumor effects against hepatocellular carcinoma"

### Supplementary Materials

#### Supplementary Materials and Methods

#### Cells

PBMCs were from local Shepeard Community Blood Center (Augusta, GA) and human T cells were isolated by EasySep human T cell isolation kit (Stemcell, Vancouver). The Jurkat, HEK293T, and HepG2 cells were from ATCC (Manassas, VA). Huh7 cells were from Dr. Satyanayana Ande, Georgia Cancer Center. Cells were cultured according to ATCC recommendation, and no more than 10 passages was used to assure authenticity. HepG2-Luc and HepG2-GFP cells were created in the lab by transduction of original HepG2 (passage 3) with lentivector expressing IRFP-Luciferase or GFP. Positive cells were sorted by GFP or IRFP.

#### Mice and tumor models

BalB/C, NOD-*scid* IL2Rgamma$^{null}$ (NSG) mice were from JAX (Bar Harbor, ME). Animal protocol is approved by the IACUC of Augusta University. Subcutaneous (SC) and intraperitoneal (IP) tumors were established by inoculating the indicated number of HepG2 or its derivatives. Tumor volume was calculated by formula: ½(LxWxH), or by bioluminescent intensity (BLI) using the Ami X spectral imaging system. BLI values (total photon/s) were analyzed using Amiview software.

##### Generation of hGPC3-specific mAbs

The hGPC3-specific mAbs were generated as previously descried (1). Female BALB/c mice of age 6-10 weeks were immunized with 100µg recombinant hGPC3 protein (AA25-560) (Creative Biomart) emulsified with complete Freund's adjuvant (Sigma-Aldrich) by footpad injection. Three weeks later, mice were boosted twice with 70µg recombinant hGPC3 together with incomplete Freund's adjuvant (Sigma-Aldrich) by footpad injection at 2-week intervals. A final boost was carried out by tail vein injection of 50 µg recombinant hGPC3 in 100 µl PBS. Three days after the final booster, mice were sacrificed and splenocytes were harvested to fuse with Sp2/0 myeloma cells. Hybridomas were created from the immunized splenocytes when a high level (>1:256,000) of anti-hGPC3 Abs was detected in the serum. Hybridomas were selected by HAT medium, and the positive clones were screened by indirect ELISA coated with 5 µg/ml hGPC3. Sub-cloning was performed three times using the limiting dilution method to obtain mAb. The mAbs were further characterized by intracellular staining of hGPC3-N and hGPC3-C fragment transfected 293 cells and by immune fluorescent staining of HepG2 tumor cells. The FITC labeled anti-mouse Ig κ chain antibody was used as secondary antibody in the immune fluorescent staining as all the antibody contain κ chain. A few mAbs were harvested and purified from ascitic fluid of BALB/c mice. The concentration and isotypes of the mAbs were determined using BCA protein assay kit and a mouse mAbs isotyping kit, respectively.

##### Identification of the mAb's cDNA sequences

To obtain the cDNA sequences of mAbs, we used the technique of 5′ rapid amplification of complementary DNA ends (5' RACE) (2, 3). Briefly, total RNA was isolated from hybridomas, and complementary DNA was generated with an oligo dT primer. PolyC was added to the 5′ end of complementary DNA by terminal transferase. Then, PCR was conducted to amplify the cDNA by using the 5′ pGI primer: CACCGGGIIGGGIIGGGIIGG, and one of the following 3′ primers corresponding to the constant (C) region of the mouse IgM heavy chain or the kappa chain:

1. For 6G11 and 12D7, IgM heavy chain primer (mIgM-OP) and κ chain primer (mIgk-OP) were designed as 3’ primers.
    a. Mouse IgM heavy chain primer (mIgM-OP): CCTGGATGACTTCAGTGTTGTTCTG;
    b. Mouse κ chain primer (mIgk-OP): CACTGCCATCAATCTTCCACTTG.
2. The PCR products were purified and sequenced. Based on the preliminary sequences, internal 3’ primers were designed. Together with 5’ pGI primer, a second and more specific PCR was run.
    a. Mouse IgM heavy chain internal 3’ primer (mIgM-IP): GGGGGAAGACATTTGGGAAGG;
    b. Mouse κ chain internal 3’ primer (mIgk-IP): CACTGGATGGTGGGAAGATGG.
3. Because the immunoglobulin class of 8F8 is IgG2a, different heavy chain 3’ primers were used:
    a. Mouse IgG2a heavy chain primer (mIgG2aH-OP): GGTTACAGTCACTGAGCTGCTG;
    b. Mouse IgG2a heavy chain internal primer (mIgG2aH-IP): GGAGCCAGTTGTATCTCCACAC

**Generation of recombinant antibody**

The cDNA sequences of mAb 6G11, 8F8, and 12D7 were used to generate recombinant mAb. Briefly, the recombinant mAbs were constructed by fusing their VH or VL with the constant region (CL or CH) of human Igγ1 or Igκ (4) (Fig. 3A). The L chain and H chain genes were synthesized and placed under the CMV promoter and co-transfected into 293 cells. The mAbs were purified with Protein-G column. SDS-PAGE was performed to determine the purification and molecular weights of mAbs. Thirty-three pmol (5 μg) of each antibody in either non-reducing or reducing condition were added, and the samples were run on 4-12% polyacrylamide gels with 4% stacking gels (Invitrogen). The gel was then stained using the SimplyBlue SafeStain (Invitrogen).

**Epitope mapping**

The epitopes recognized by mAbs were mapped as follow:

1. 1st, we determine whether the mAb recognize hGPC3 N (25-358aa) or C (359-560aa) fragments. Both Western Blot analysis and intracellular staining were used. Through the 1st step, we identified that 6G11 and 12D7 bound to hGPC3-N fragment, while 8F8 bound to hGPC3-C fragment.
2. 2nd, to map the epitopes in hGPC3-N fragment, we further divided the N fragments into 3 shorter fragments (hGPC3-N1, 1-126aa; hGPC3-N2 (118-242aa; and hGPC3-N3, 232-358aa) and expressed them in 293T cells. Then, the transfected 293T cells were staining with the corresponding mAbs to determine whether which fragment the mAbs would bind. By doing so, we determined that 6G11 bound to the 25-126aa fragment. However, we were unable to determine the epitope of 12D7.
3. 3rd, we divided the 25-126aa fragment into shorter peptides:
    1) Pep-N1: QPPPPPPDATCHQVR (25-39);
    2) Pep-N2: PPDATCHQVRSFFQRLQPGLKWVPETPVPGSDLQVCLPK (30-68);
    3) Pep-N3: GSDLQVCLPKGPTCCSRKMEEKYQLTARLNMEQLLQSAS (59-97);
    4) Pep-N4: NMEQLLQSASMELKFLIIQNAAVFQEAFEIVVRHAKNYT (88-126).
4. To map the epitope in the C fragment (359-560aa) recognized by 8F8 mAb, we used the following peptides:
    1) C1: GKLCAHSQQRQYRSAYYPEDLFIDKKVLKVAHVEHEETL
    2) C2: AHVEHEETLSSRRRELIQKLKSFISFYSALPGYICSHS
    3) C3: LPGYICSHSPVAENDTLCWNGQELVERYSQKAARNGMK
    4) C4: QKAARNGMKNQFNLHELKMKGPEPVVSQIIDKLKHINQ
    5) C5: IDKLKHINQLLRTMSMPKGRVLDKNLDEEGFESGDCGD
    6) C6: GFESGDCGDDEDECIGGSGDGMIKVKNQLRFLAELAYD
    7) C7: RFLAELAYD LDVDDAPGNSQQATPKDNEISTFHNLGNVH
5. ELISA assay with synthetic peptides

    96-well Flat-Bottom Immuno Plates (from Thermo Fisher) were coated with 4ug of PBS-diluted peptides, overnight, at 4°C. 1ug of rhGPC3 (from Creative Biomart) was used as positive control. The next day, peptides were discarded and plates were blocked with 100µl of 5% non-fat milk in PBS. Then, the plates were washed with 300µl/well of PBST (0.1% Tween20). Next, incubation with 100uL/well of different antibody dilutions. This was followed by four washes with PBST. Then, incubation with 100uL/well of anti-human IgG-HRP (From KPL) (1:2000 dilution). This was followed by four washes with PBST. Next, 100uL/well of TMB (from Biolengend), incubation for 30 minutes. Then, 100uL/well of 2N $H_2SO_4$. Lastly, OD450nm was detected in an xMark™ Microplate Spectrophotometer (from BIO-RAD). If not indicated otherwise, incubations were performed for 1 hour at RT, and antibodies were diluted in 2% non-fat milk in PBS.

**Measurement of mAb affinity by BioLayer Interferometry**

The affinities of recombinant mAbs to hGPC3 were tested using Biolayer interferometry as previously described (5). The recombinant mAbs were diluted to the concentration of 5 µg/mL with PBS containing 0.05% Tween 20 and immobilized onto Anti-hIgG Fc Capture (AHC) biosensors (Sartorius AG). Following a 60-seconds baseline step with PBST, biosensor tips were immersed into the wells containing serial dilutions of hGPC3 antigen. After an association of 300 seconds, a dissociation step of 600 seconds was conducted. The $K_D$ were calculated using a 1:1 binding model in Data Analysis Software 9.0.

#### Generation of CARTs

The second generation of CARs was constructed as follow. The CAR includes the leader sequence of human CD8 molecule, the VL and VH, a glycine and serine (GS) linker between the VL and VH, a fragment of CD8 hinge, the transmembrane domain of human CD28, and the 4-1BB and CD3 ζ chain signaling domains. In some cases, the mCherry is fused to C-terminal of CD3ζ chain to tag the CAR. The entire expression cassette was inserted behind the EF1α promoter in the lentivector (lv). Lentivector production was as described previously (6) by co-transfection of 293T cells with 4 plasmid DNA including the shuttle CAR expression shuttle plasmid, pLP1, pLP2 and pVSVG (Invitrogen). Virus particles in the media were concentrated by high speed centrifugation and sucrose gradient as previously described (6, 7). Virus titer was determined by transducing the Jurkat cells with a serial dilution of virus stock and anti-mouse Fab antibody staining (Jackson ImmunoReseach Lab). Human primary T cells were isolated transduced by lentivectors at 20 MOI as previously described (3). Details are in Supplementary Materials and Methods

#### Flow cytometry analysis

Cells from blood, spleen, or tumors were stained with the indicated antibodies and analyzed on flow cytometers of LSR (BD Bioscience) or NovoCyte Quanteon (Agilent). Zombie green was added to exclude dead cells in certain staining. The antibodies included Biolegend CD45, CD8, CD4, PD1, LAG-3, Annexin-V, IFNγ, and IL2. The goat anti-mouse Fab antibody was from Jackson ImmunoReearch. The FITC-labeled hGPC3 was from Acro Biosystems.

#### Intracellular staining of cytokines

To study the cytokine production by the CARTs in the tumor lesion, single cells from tumor lesions were co-cultured with $5x10^5$ HepG2 tumor cells according to the % of CAR+ T cells in the single cell suspension, and desired E/T ratios. Monensin was added to the co-culture for the last 4h before collecting the cells for intracellular staining of the IL2 and IFNγ.

#### ELISA and lactate dehydrogenase (LDH) cytotoxicity assay

HepG2 cells ($1\times10^4$/well) were plated on 96 round-well plates and grew overnight. CARTs were added at indicated effector/target (E/T) ratios. After 20hrs co-culture, the cytotoxicity was measured by the LDH assay according to manufactory's protocol (Promega). The IFNγ production was determined by ELISA (Biolegend).

**Real time cell analysis of cytotoxicity of CARTs**

$1\times10^4$ HepG2-GFP$^+$ cells were plated in duplicate on flat-bottom 96 well plate. After overnight, CARTs at indicated E/T ratio was added and co-cultured up to 72 h. Plates were imaged every 3h using the IncuCyte SX1 Live-Cell analysis system (Sartorius). Five images per well were collected at each time point. Total integrated GFP area was assessed at each time point to measure the live, GFP$^+$ tumor cells using Sartorius software and plotted as kinetics of time.

**Luminex assay**

Simultaneous quantification of cytokines in supernatants was performed using LEGENDplex Human CD8/NK Panel (13-plex) with V-bottom Plate (BioLegend Cat# 740267) according to manufacturer's instructions. In brief, samples were thawed completely, mixed, and centrifuged to remove particulates prior to use. Supernatants were diluted with Assay Buffer and standards were diluted with culture media accordingly. Standards and samples were plated with capture beads for TNF-α, IFN-γ, IL-4, IL-2, IL-6, IL-10, IL-17A, sFas, sFasL, Granz. A, Granz. B, Perforin, Granulysin and incubated for 2 h at room temperature on plate shaker (800 rpm). After washing the plate with Wash Buffer, Detection Antibodies were added to each well. Plate was incubated on shaker for 1h at room temperature. Finally, without washing, SA-PE was added and incubated for 30 min. Samples were acquired on CytoFLEX flow cytometer (Beckman Coulter Life Sciences). Standard curves and protein concentration were calculated using R package DrLumi (Sanz et al., 2017) installed on R 3.5.2 (https://www.r-project.org/). Limit of detection was calculated as average of background samples plus 2.5 x SD. Assay and data calculations were performed in Immune Monitoring Shared Resource (Augusta University).

**Immunohistochemical (IHC) Staining and scoring**

The paraformaldehyde-fixed and paraffin-embedded blocks of health liver, adjacent normal liver and HCC tissue are from cases operated at Lombardi Cancer Center of Georgetown University. Tissues were sectioned and stained in the clinical pathology lab of Georgetown cancer center. All patients gave informed consent, and the study was authorized by the respective Hospital Ethics Committees. Pathologic slides were stained with one of the following purified mAbs (6G11, 12D7, and 12E6, 8C8 and 8F8). Secondary antibodies were either goat-anti-mouse IgM or goat-anti-mouse IgG (Vector Labs). Immunoscores were calculated by adding intensity and region of staining: 0, 1, 2, 3 correspond to no, weak, moderate, and high intensity staining; 0, 1, 2, 3 correspond to no, focal as less than 10% of cells, regional as 10%–50% of cells, and diffuse region as >50% of cells staining. To detect tumor infiltrating T cells by IHC, tumor tissues were collected and fixed. The paraffin embedded tissues were sectioned and stained for human CD8 T cells by anti-CD8 antibody (Sino Biologicals, Cat#: 10980-T24) in Augusta University Histology core lab.

**Tumor Infiltrating Lymphocytes (TILs):**

For ex vivo analysis of CARTs in the tumor lesions, mice were euthanized, solid tumor tissue was collected, washed in PBS, mechanically dissociated and digested with digestion medium (RPMI supplemented with 0.1% collagenase type V (Sigma C9263), 0.1% hyaluronidase (Sigma H6254), and 100 unit/ml DNase I (Sigma D4527)) for 30-45 minutes at 37$^{o}$C. After RBCs lysis, the single cell suspension was ready for cytotoxicity assay by IncuCyte real time analyzer.

**Bulk RNA Sequencing.** The RNA-seq libraries were prepared from total RNA and sequenced by Novogene. The paired-end 150bp raw sequencing reads were examined by FastQC v 0.11.8. Adaptor sequences and low-quality bases were trimmed using Trim Galore! v 0.6.3. The cleaned reads were mapped to reference genome (mm10) using STAR aligner v 2.7.8a. The QC and mapping steps were performed in Galaxy server (use.galaxy.org) The sequence counts for each gene were collected by Rsubread featureCounts functions v 1.22.2. The differential gene expression analysis was performed using DESeq2 v1.32.0. The heatmaps were generated using ComplexHeatmap v2.8.0 in R v4.1.

#### Statistical analysis

Statistical analyses were performed using student two-tail *t*-test or 2way ANOVA in the Prism software (GraphPad Inc.) or special software indicated on the figures.

##### References cited in Supplementary Materials and Methods


1. Xiao H, Zhu P, Liu B, Pan Q, Jiang X, Xu X, Fu N. 2010. Generation and characterization of human delta-globin-specific monoclonal antibodies. *Blood Cells Mol Dis* 44: 127-32
2. Walchli S, Loset GA, Kumari S, Johansen JN, Yang W, Sandlie I, Olweus J. 2011. A practical approach to T-cell receptor cloning and expression. *PLoS One* 6: e27930
3. Zhu W, Peng Y, Wang L, Hong Y, Jiang X, Li Q, Liu H, Huang L, Wu J, Celis E, Merchen T, Kruse E, He Y. 2018. Identification of alpha-fetoprotein-specific T-cell receptors for hepatocellular carcinoma immunotherapy. *Hepatology* 68: 574-89
4. Tiller T, Meffre E, Yurasov S, Tsuiji M, Nussenzweig MC, Wardemann H. 2008. Efficient generation of monoclonal antibodies from single human B cells by single cell RT-PCR and expression vector cloning. *J Immunol Methods* 329: 112-24
5. Chi X, Yan R, Zhang J, Zhang G, Zhang Y, Hao M, Zhang Z, Fan P, Dong Y, Yang Y, Chen Z, Guo Y, Zhang J, Li Y, Song X, Chen Y, Xia L, Fu L, Hou L, Xu J, Yu C, Li J, Zhou Q, Chen W. 2020. A neutralizing human antibody binds to the N-terminal domain of the Spike protein of SARS-CoV-2. *Science* 369: 650-5
6. He Y, Zhang J, Mi Z, Robbins P, Falo LD, Jr. 2005. Immunization with lentiviral vector-transduced dendritic cells induces strong and long-lasting T cell responses and therapeutic immunity. *J Immunol* 174: 3808-17
7. Liu Y, Peng Y, Mi M, Guevara-Patino J, Munn DH, Fu N, He Y. 2009. Lentivector immunization stimulates potent CD8 T cell responses against melanoma self-antigen tyrosinase-related protein 1 and generates antitumor immunity in mice. *J Immunol* 182: 5960-9.

#### Supplementary Figures (13)

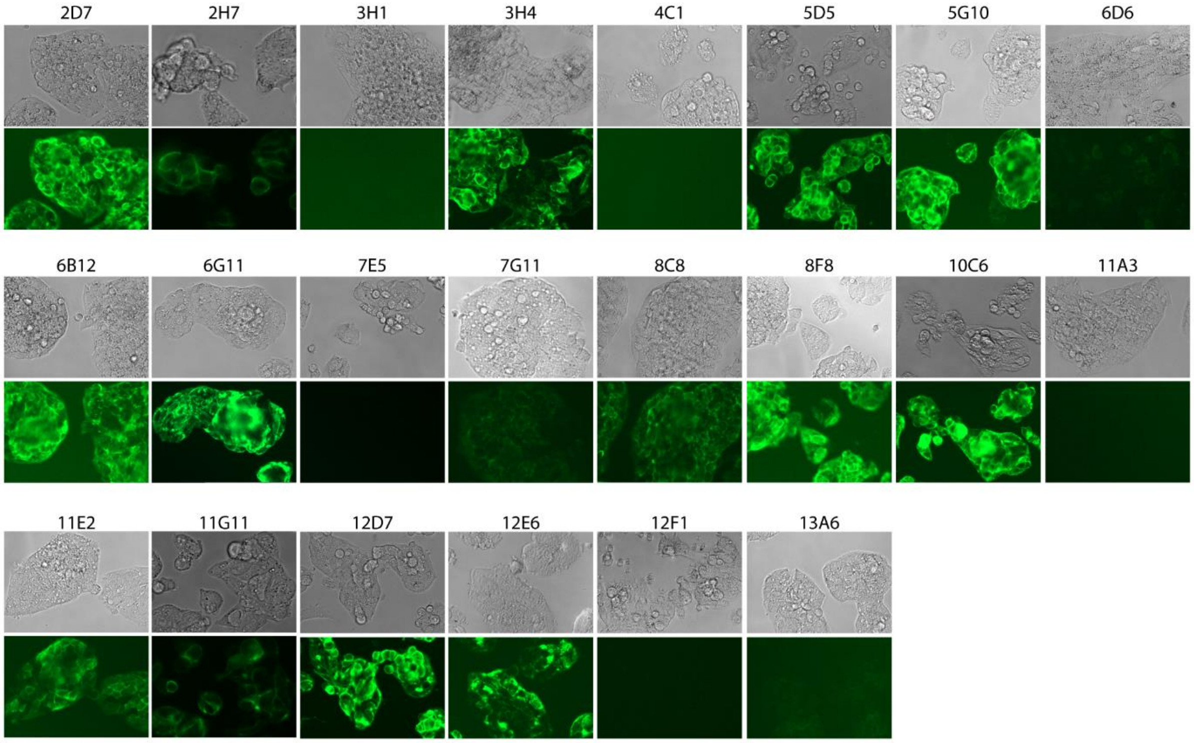


fig S1. New hGPC3-specific mAbs bind HepG2 tumor cells. HepG2 tumor cells were stained with the indicated mAbs and then with FITC labeled anti-mouse kappa chain antibody. Pictures were taken in Phase and Green channel.

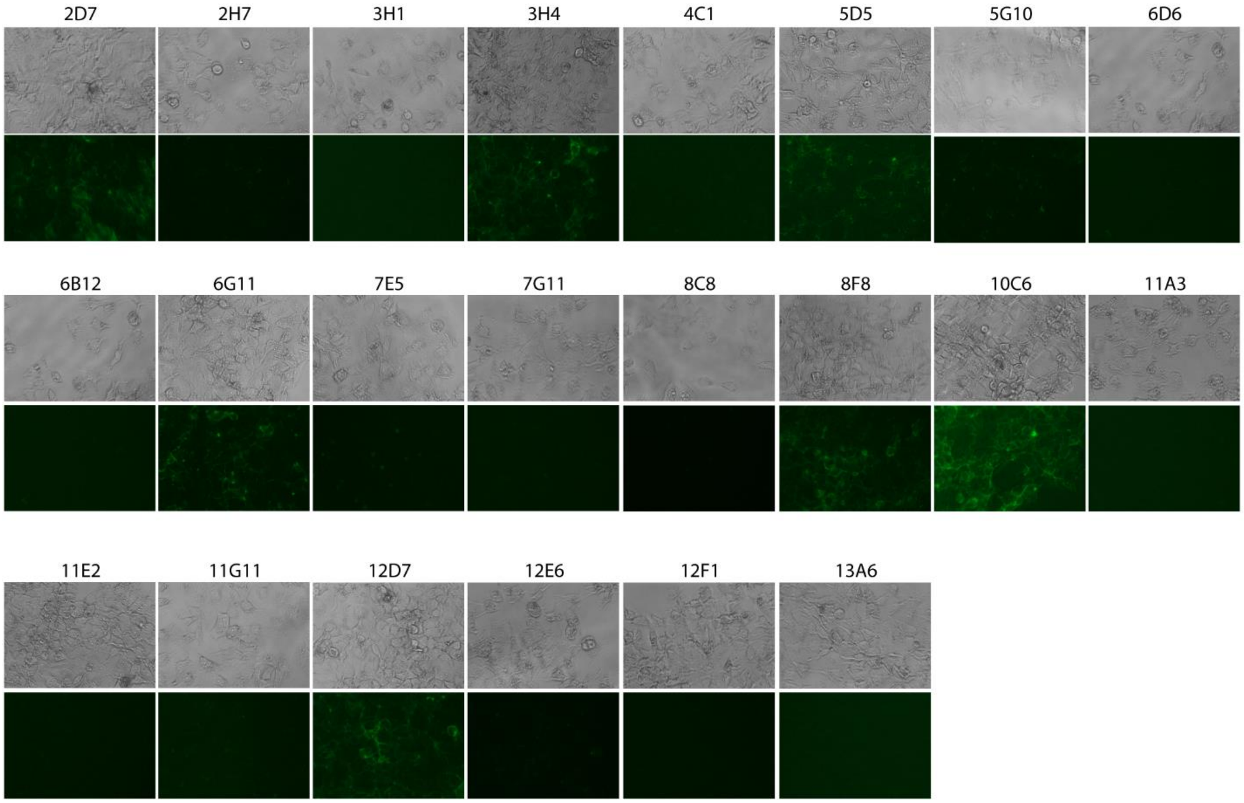


fig S2. New hGPC3 mAbs stain hGPC3low Huh7 tumor cells. Similar to fig. S1, Huh7 tumor cells were stained with mAbs and then FITC labeled secondary anti-mouse kappa chain antibody. Seven mAbs (2D7, 3H4, 5D5, 6G11, 8F8, 10C6, and 12D7) are able to stain Huh7 tumor cells.

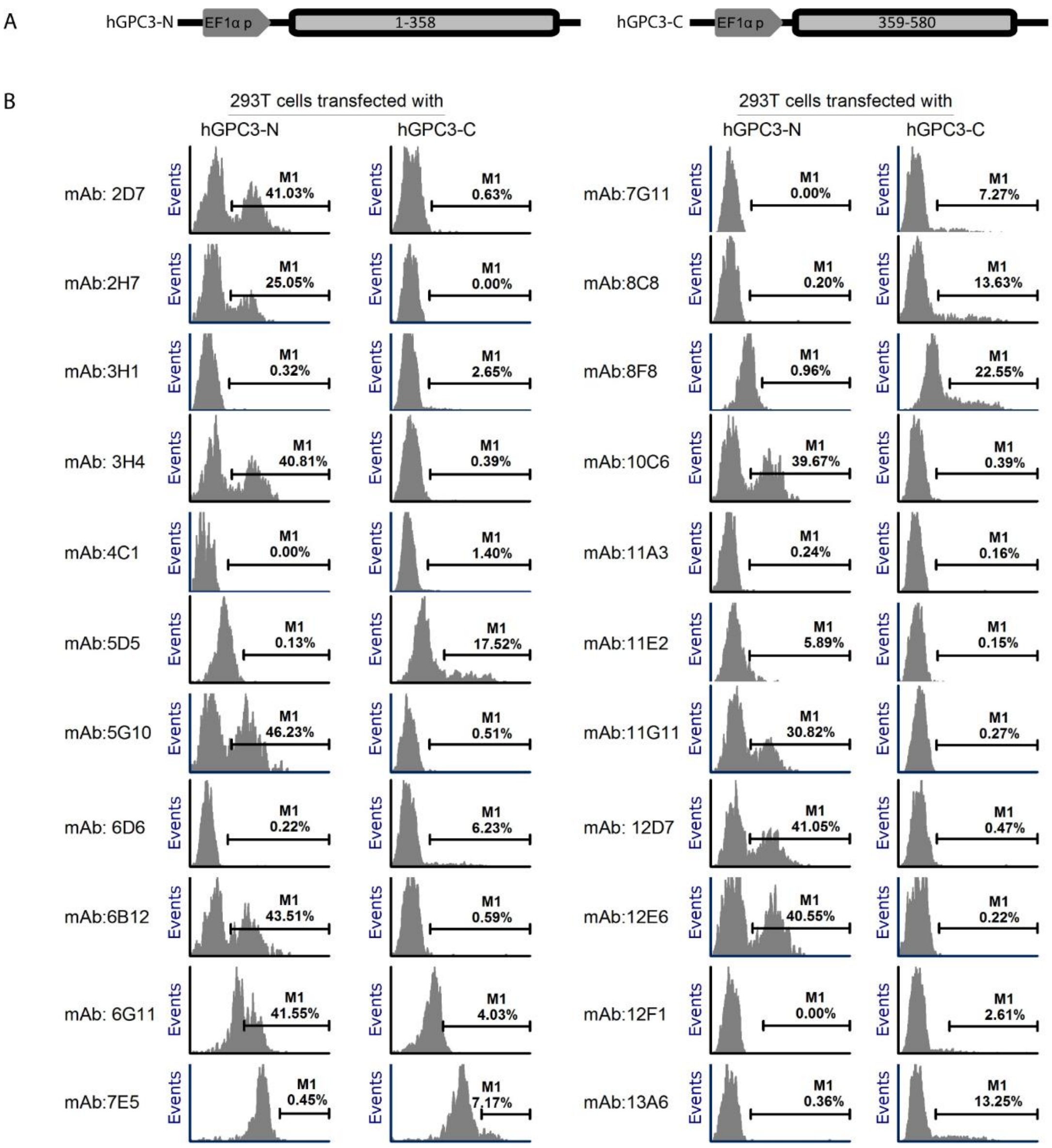


fig S3. The mAbs binding to hGPC3-N and hGPC3-C fragment are identified. (A) The plasmid DNA expressing the N- fragment (1-358aa) and C- fragment (359-580aa) were presented. Initiation codon was added to the C-fragment. (B) Intracellular staining of the 293T cells transfected with hGPC3-N or hGPC3-C by different hGPC3-specific mAbs. The gate was set by the parental 293 cells.

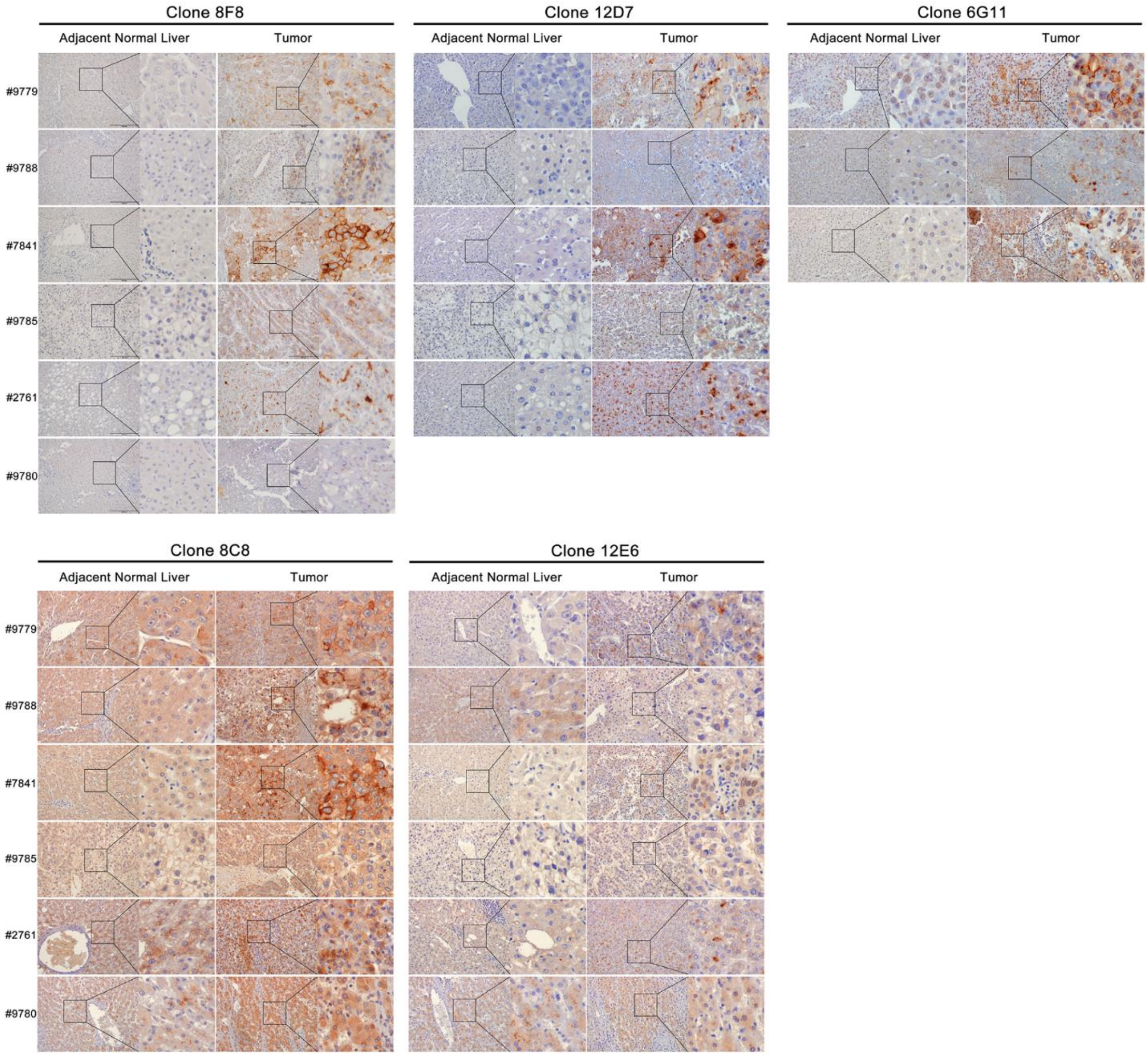


fig. S4. The 6G11, 8F8, and 12D7 mAbs stain only the HCC tumor tissues. IHC staining of HCC tumor tissues and their paired adjacent normal liver tissues by the 5 indicated mAbs. HRP-conjguated anti-kappa chain antibody was used as the secondary antibody.

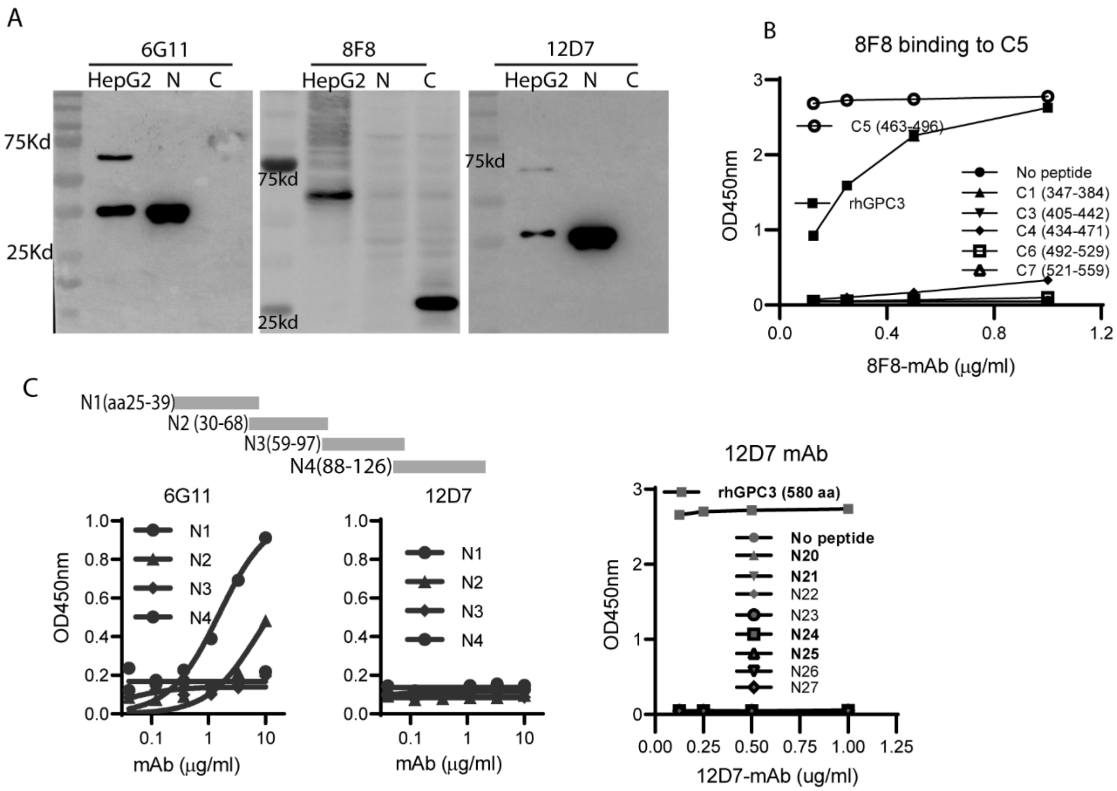


fig S5. 6G11, 8F8, and 12D7 mAbs bind to distinct epitopes on hGPC3. (A) Western Blot analysis of the 293T cells transfected with hGPC3 N- or hGPC3 C- fragment by indicated hGPC3-specific mAbs. The data confirmed that 6G11 and 12D7 bound to hGPC3 N-fragment and 8F8 bound to the hGPC3 C-fragment. HepG2 cell lysate was used as positive control. (B and C) Mapping the binding epitopes of 6G11, 8F8, and 12D7 by overlapping peptides. The data showed that 8F8 bound to C5 peptide (AA463-496) (B) and 6G11 mAb bound to N1 peptide (AA25-39) (C). 12D7 bound no specific epitope. The N20 to N27 peptides cover the sequences of hGPC3 from AA120 to AA360.

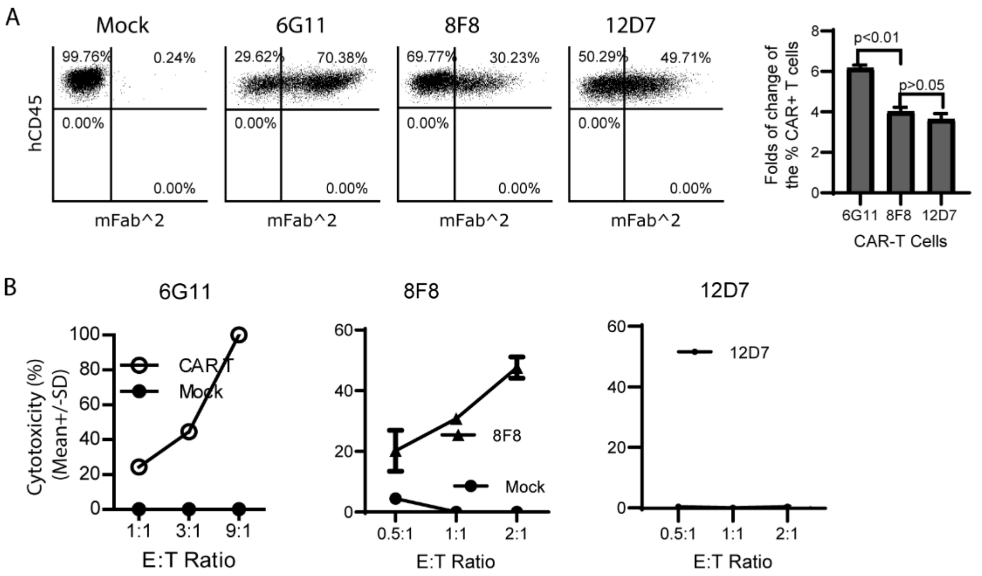


fig S6. 6G11, 8F8, and 12D7 CAR-Ts are expanded by, but only 6G11 and 8F8 CAR-Ts kill hGPC3+ tumor cells. (A) Mock-T and CARTs were co-cultured with HepG2 tumor cells. Representive dot plots after stimulation by HepG2 cells were shown. The % of CAR+ T cells were compared to CARTs before co-culture (Fig. 3C) and the fold of increase of CAR+ T cells were calculated. (B). The cytotoxicity of CARTs on Huh7 tumor cells. (B) The 6G11 and 8F8 CARTs , but not the 12D7 CARTs could kill the $hGPC3^{low}$ Huh7 tumor cells. Each dot represents an average of triplicate data. The experiments were repeated 3 times with similar observation.

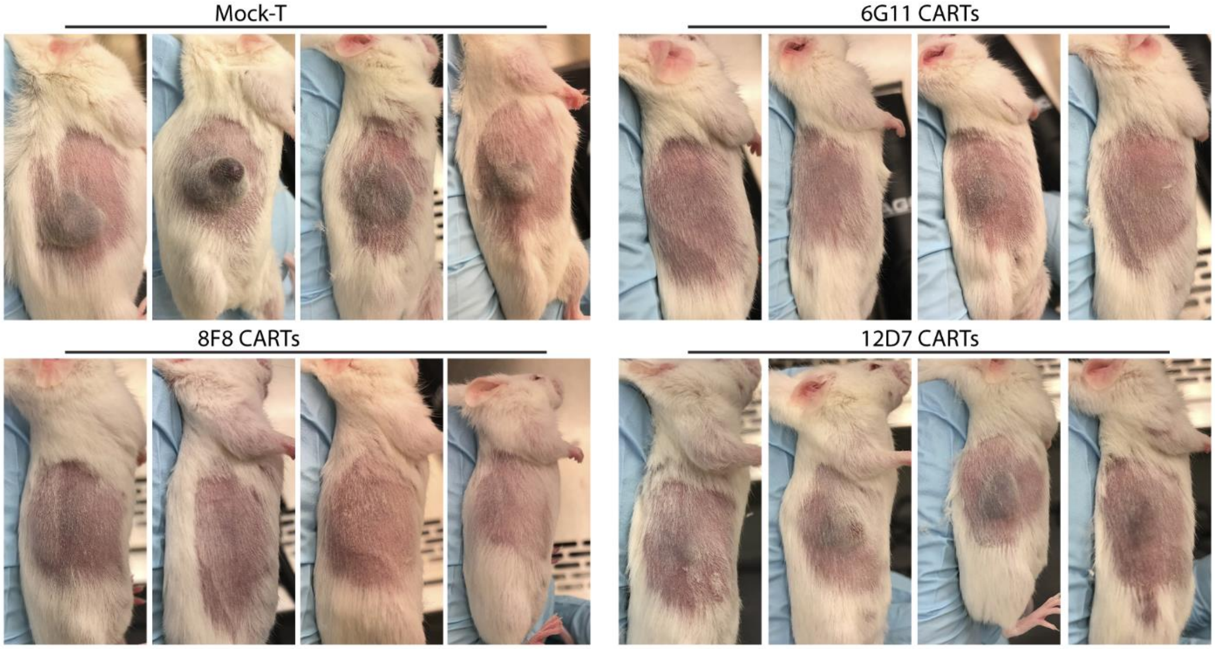


fig S7. 6G11 and 8F8 CARTs generate potent antitumor effects. Experiment was done as depicted in Fig 3. The mouse tumor pictures were taken at 40 days after tumor inoculation (33 days after treatment). Representative tumor pictures from one of the 5 experiments were shown.

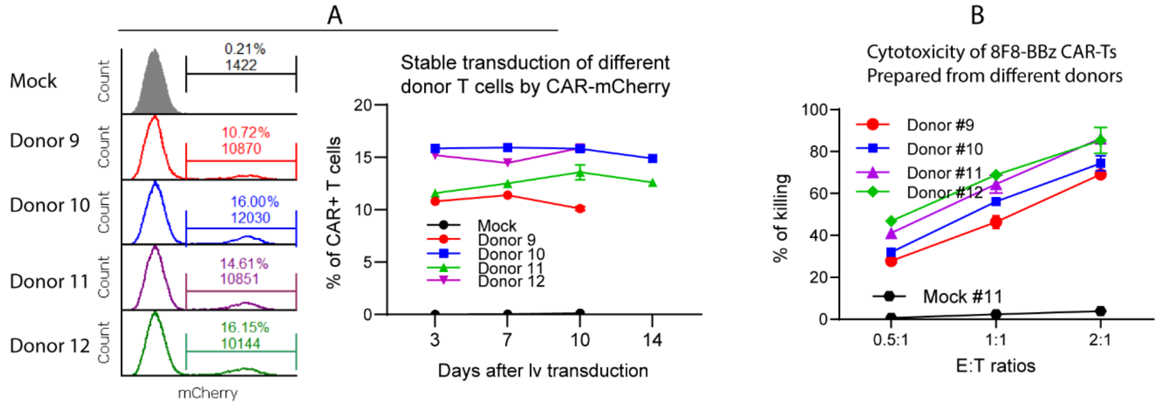


fig S8. Multiple donor T cells can be comparably transduced by 8F8-BBz CAR with mCherry tag. (A). Primary human T cells from 4 donors were activated and transduced by 8F8-BBz CAR with mCherry tag at MOI 20. The mCherry+ CARTs were monitored for 2 weeks. (B). The cytotoxicity of 8F8-BBz CARTs from 4 donors were determined on D10 after transduction.

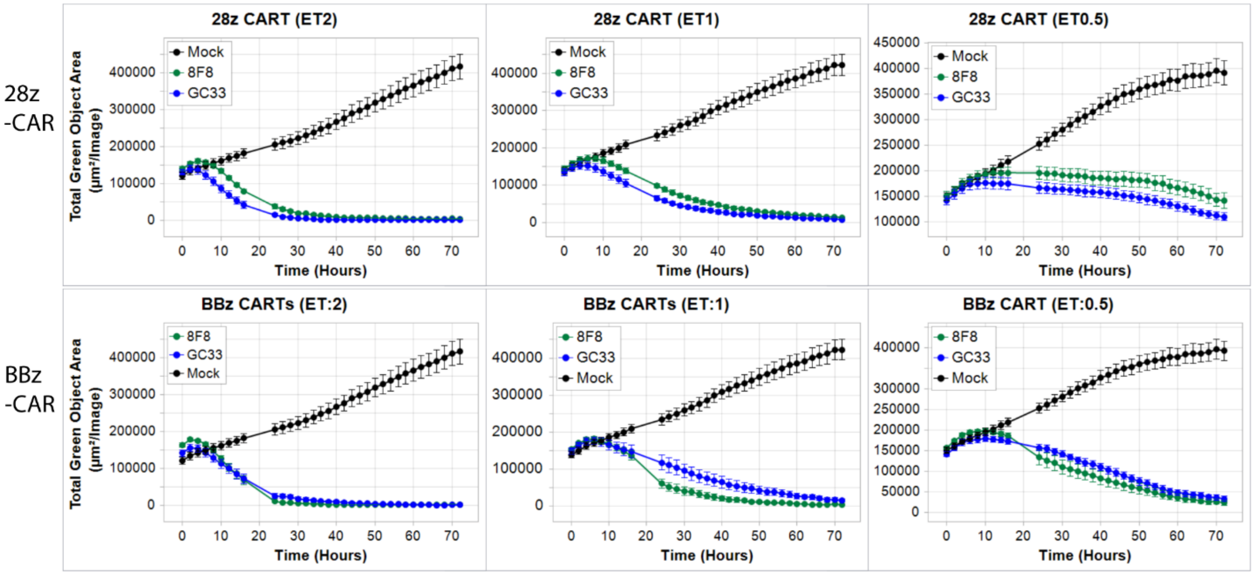


fig S9. Real-time cell analyzer showed 8F8 and GC33 CARTs have similar cytotoxicity. Analysis of CTL activity of different CARTs by IncuCyte real-time cell analyzer. 10,000 HepG2-GFP cells were plated onto flat 96 well plate overnight, the next day, CARTs were added at indicated ET ratios and let the cells co-cultured for 3 days. Every 3hours, 5 picture were taken for each well. Each data point represents 10 pictures from 2 duplicate wells. The loss of GFP+ tumor cells was recorded by photos and analyzed.

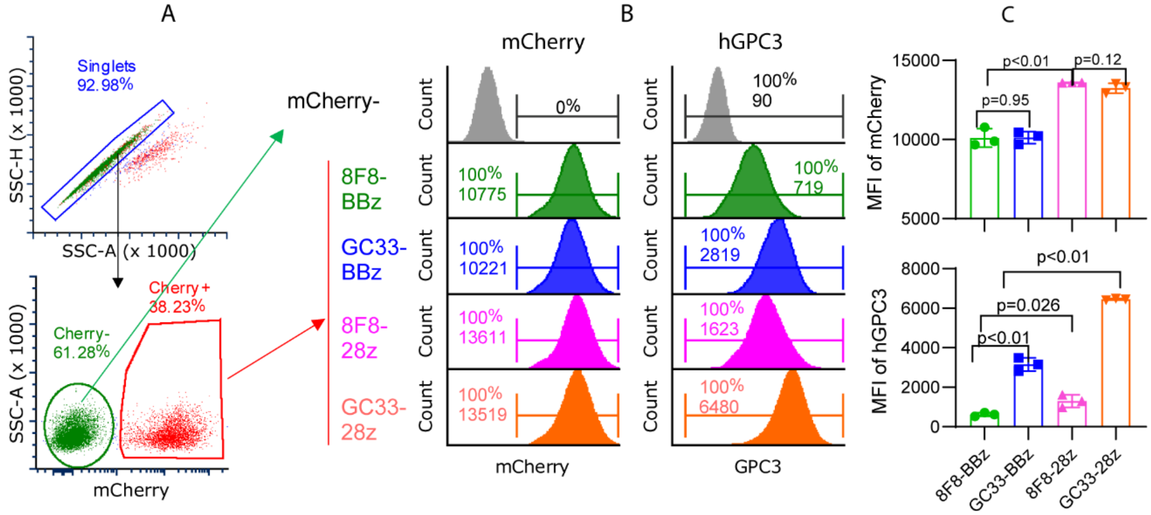


fig S10. 8F8 CARTs have lower avidity of binding soluble hGPC3 protein than GC33 CARTs. T cells were transduced with 8F8 or GC33 BBz/28z CARs. On Day 10 after transduction, cells were stained with FITC labeled hGPC3 protein. (A and B) Shown are the representative dot plots and histogram of mCherry and hGPC3 on CARTs. The mCherry+ T cells were gated as CAR+ T cells. mCherry negative T cells were gated as control. (C). A summary of data from 3 time points (D10, D14, and D17 after lv transduction) was shown. By mCherry, the CAR expression level is similar between 8F8 and GC33, but 28z CAR level is 1.3 times higher than BBz CAR. The 8F8 CAR binds 4x less of hGPC3 protein than GC33 CAR, and the 28z CAR of both 8F8 and GC33 bind twice more of hGPC3 than their BBz CAR counterparts.

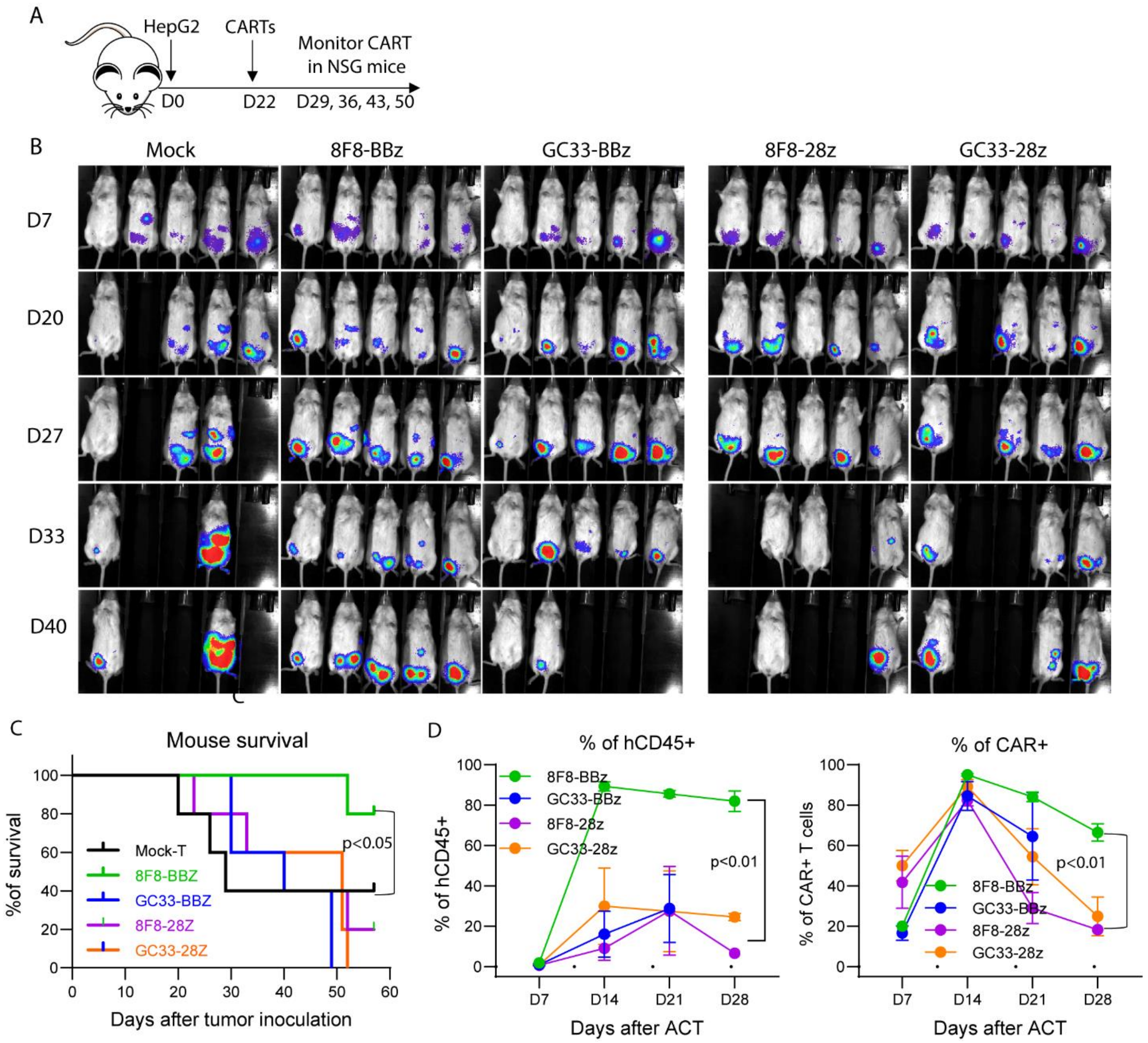


fig S11. 8F8-BBz CART generates enhanced expansion and extends the survival of mice bearing large IP tumors. (A) The experimental scheme is shown. (B) Bioluminescent imaging of IP tumors at the indicated day after tumor inoculation. (C) Survival of IP tumor bearing mice among different groups. (D) The % of hCD45 and CAR+ T cells in mouse blood. The gating strategy is as of Fig. 5D. Statistical analysis was done as described in Fig. 5.

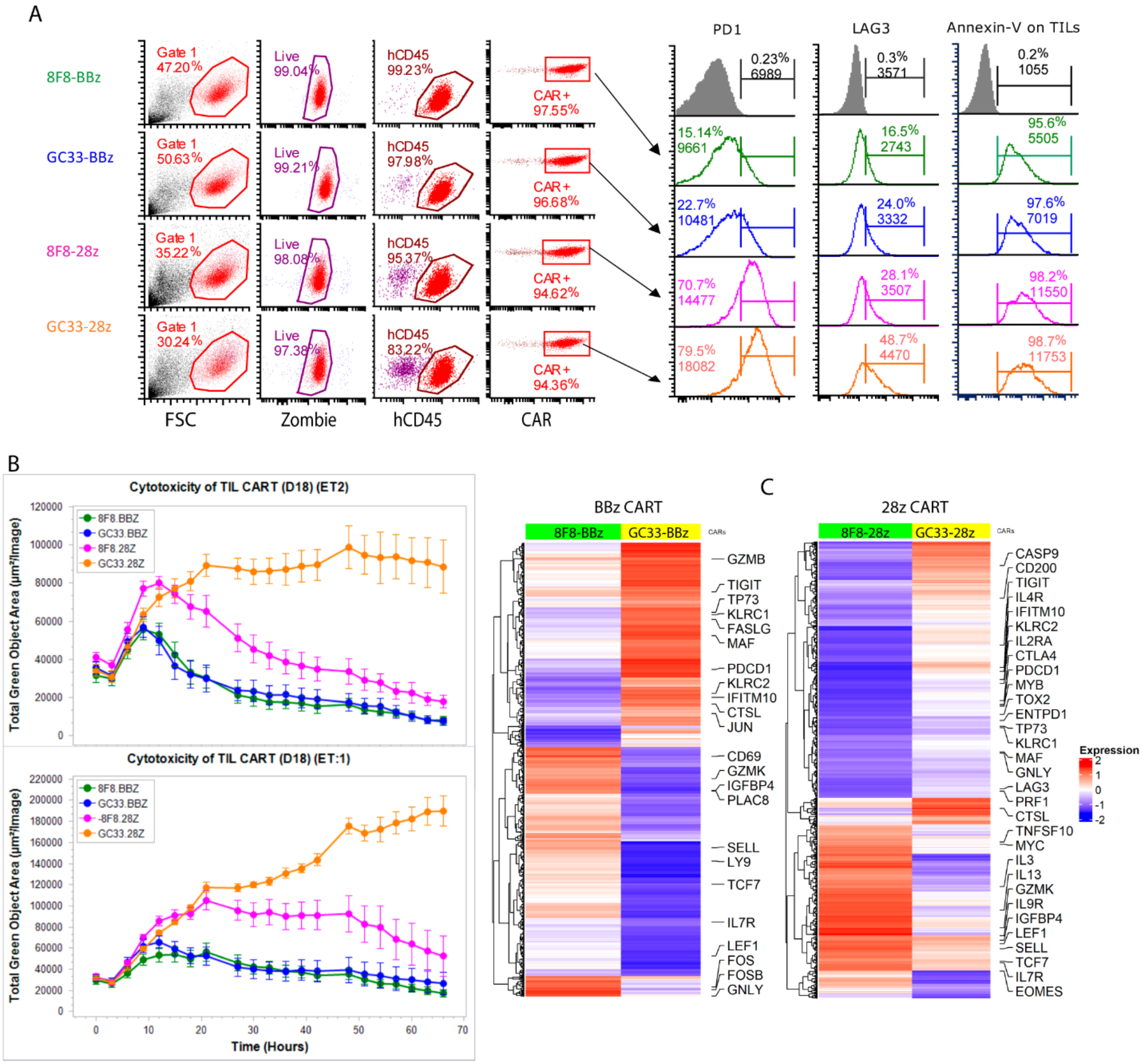


fig S12. 8F8-BBz CARTs are less exhausted and more functional in the tumors at day 18 after ACT. (A) Single cell suspensions were prepared from each tumor and cells were stained for hCD45 and exhaustion markers or apoptosis. (B) Cytotoxicity of the tumor infiltrating CAR-ts were analyzed by real time cell analyzer at indicated E:T ratio. The GC33-28z CART already lost most of its killing activity. (C) The differentially expressed genes between 8F8 and GC33 CART cells showed that 8F8 CARTs were less exhausted and more functional than GC33 CARTs.

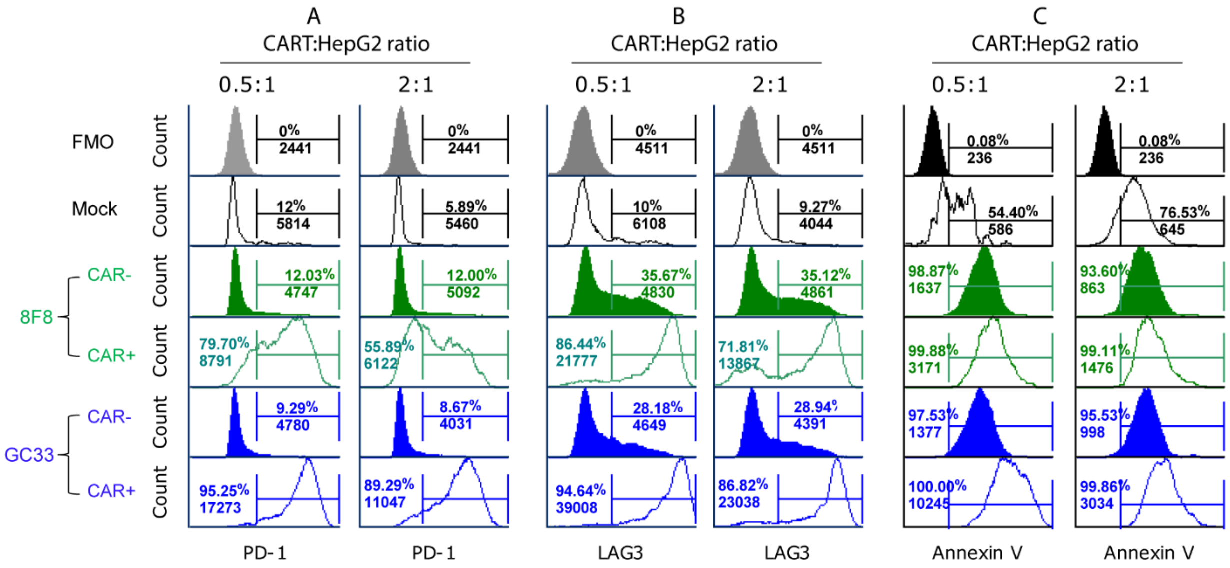


fig S13. 8F8-28z CART is less exhausted and apoptotic than GC33-28z CART after co-culture with tumor cells. CART and HepG2 tumor cells were co-cultured for 2 days at indicated ratios, apoptosis and exhaustion markers were stained. The gating strategy is the same as Fig.7. (A and B). Exhuation marker of PD1 an LAG3 were presented on CAR- and CAR+ T cells. FMO: fluorescent minus one (PD1 or LAG3). (C). Apoptosis on CAR- and CAR+ T cells were shown.
